# Paracrine Factor Local Gradient-Generating System for Engineering Perfusable Vascularized Hepatocyte Tissues with Perfusion-Induced Proliferation

**DOI:** 10.1101/2025.11.10.687539

**Authors:** Yen-Hsiang Huang, Tadahiro Yamashita, Ryo Sudo

## Abstract

Donor organ shortages drive the urgent need for engineered hepatocyte tissues, yet functional vascular integration remains a major bottleneck in liver tissue engineering. Current vascularization strategies struggle to achieve perfusable microvessels that penetrate hepatocyte tissue. Furthermore, in vitro recapitulation of hepatic regeneration remains a major challenge. This study presents a paracrine factor local gradient (PFLG)-generating system that constructs vascularized, perfusable hepatocyte tissues and recapitulates the perfusion-mediated proliferative capacity of primary hepatocytes. The PFLG-generating platform integrates fibroblast-loaded cryogels with a microfluidic device to direct angiogenesis prior to hepatocyte seeding, thereby enabling microvessels to penetrate three-dimensional hepatocyte tissue. Within the vascularized constructs, microvessels directly penetrated the hepatocyte parenchyma, recapitulating intimate hepatocyte–microvessel contact in vivo. These constructs enhanced hepatocyte polarity and functional bile canaliculi. Importantly, under perfusion culture, robust hepatocyte proliferation was induced, with Ki67-positive hepatocytes increasing significantly, including mitotic cells while preserving polarity. By contrast, this proliferative response was minimal under static conditions. Time-lapse imaging and functional assays confirmed perfusion through penetrating microvessels. These findings demonstrate that perfusion-mediated cues are essential for inducing hepatocyte proliferation while maintaining functional polarity. This modular and programmable culture platform lays the foundation for scale-up toward transplantable liver tissues.

## Introduction

Persistent donor-organ shortages constrain liver transplantation, thereby driving the in vitro development of engineered hepatic tissues. However, in conventional culture systems primary hepatocytes rapidly lose differentiated functions and proliferative capacity, posing a major challenge for fabricating functional hepatic tissues. Clinically viable hepatic constructs must satisfy three key requirements: (i) sustained hepatocyte differentiated functions and polarity; (ii) faithful recapitulation of the in vivo sinusoidal architecture to enable hepatocyte–endothelial interactions and efficient substance exchange;^[1,2]^ and (iii) sufficient proliferative capacity to achieve a transplantable tissue size.^[3–5]^ Current strategies rarely meet all three requirements, thereby limiting clinical translation.

Hepatocytes exhibit robust regenerative capacity in vivo, restoring liver mass within approximately 3 weeks after partial hepatectomy.^[6]^ By contrast, this regenerative potential is not recapitulated in three-dimensional (3D) in vitro culture systems, where hepatocytes largely remain quiescent.^[7]^ Following partial hepatectomy, increased portal blood flow elevates shear stress and activates sinusoidal endothelial cells (SECs), which in turn secrete hepatocyte growth factor (HGF) and Wnt ligands.^[8–11]^ HGF–c-Met signaling activates the PI3K/Akt and MAPK pathways to drive proliferation, whereas Wnt–Frizzled signaling activates β-catenin and promotes hepatocyte cell-cycle re-entry.^[6,12–14]^ Although at least these signaling pathways are involved in liver regeneration, the complete mechanism remains incompletely understood. Despite this mechanistic complexity, Chhabra et al.^[15]^ developed a culture system that triggered one facet of regenerative signaling: hepatocyte–fibroblast spheroids embedded in fibrin hydrogel and cocultured with HUVEC-lined hydrogel channels. Oscillatory flow combined with interleukin-1β (IL-1β) stimulated HUVECs to secrete prostaglandin E2 (PGE_2_), inducing hepatocyte cell-cycle entry. However, because the spheroids lacked microvessel penetration and a sinusoid-like architecture enabling direct hepatocyte– endothelial exchange, hepatocyte proliferation appeared to remain limited. These findings underscore the need to develop perfusable, vascularized hepatic tissues that recapitulate sinusoid-like architecture to elicit regenerative responses in vitro.

Numerous 3D culture strategies have been developed to maintain hepatocyte functions in vitro.^[16–24]^ However, avascular hepatocyte tissues lack hepatocyte–endothelial interactions and the physiological perfusion necessary to support hepatic functions. Vascularization is therefore essential for constructing more functional hepatic tissues. A recent study utilized microfluidic devices to construct vascularized hepatocyte tissues,^[25]^ but whether these platforms faithfully recapitulate sinusoid-like architecture remains unclear. In parallel, incorporating non-parenchymal cells as continuous sources of pro-angiogenic paracrine factors has emerged as a promising strategy for vascularization.^[26,27]^ Building on these approaches, our recent work incorporated human lung fibroblasts (hLFs) as paracrine sources of angiogenic factors within primary hepatocyte spheroids in microfluidic devices, thereby establishing a paracrine factor local gradient (PFLG) that promoted spheroid vascularization.^[28]^ Although these spheroids mimicked the hepatocyte–endothelial interface of hepatic sinusoids, excessive hLF proliferation eventually compromised microvascular perfusion. To address this limitation, strategies that deliver angiogenic cues without the direct incorporation of hLFs are required to achieve functional perfusion in vascularized hepatocyte tissues.

Here, we developed a novel PFLG-generating system that enables the formation of perfusable microvessels penetrating 3D primary hepatocyte tissues. The culture platform comprises two functional components: (i) hLF-loaded cryogels that continuously release paracrine factors and (ii) a microfluidic device that provides the spatial framework for PFLG generation. Integrating these components produces a PFLG that directs vascular extension and, ultimately, vascularization. Unlike conventional coculture strategies, this system enables pre-angiogenesis prior to hepatocyte seeding, thereby accelerating subsequent vascularization of hepatocyte tissues. First, we optimized culture conditions by examining angiogenic responses to the PFLG produced by hLF-loaded cryogels during the pre-angiogenesis phase. We then determined optimal rat primary hepatocyte (rHep):HUVEC mixing ratios to promote vascularization. Next, we characterized the resulting tissue architecture recapitulating sinusoid-like structures, assessed enhancements in hepatic function following vascularization, and confirmed functional perfusion of penetrating microvessels. Finally, we showed that perfusion-mediated stimulation enhanced hepatocyte proliferation while preserving hepatic polarity in vascularized hepatocyte tissues. Collectively, these findings provide a framework for constructing perfusable hepatocyte tissues and lay the groundwork for developing transplantable liver tissue in vitro.

## Results

### Characterization of hLF culture within gelatin cryogels

We recently demonstrated that hLFs incorporated into hepatocyte tissue can promote vascularization;^[28]^ however, hLF overgrowth might impair functional perfusion. To address this issue, we developed a novel system that generates a PFLG derived from hLFs to construct perfusable hepatocyte tissue without directly incorporating hLFs into the vascularized tissue. This system employs a gelatin cryogel loaded with hLFs (**Figure 1A**). The hLF-loaded cryogel is placed in a reservoir located behind the hepatocyte tissue, opposite the HUVEC channel (Figure 1B). This spatial configuration enables the formation of an hLF-derived PFLG across hepatocyte tissue, thereby promoting its vascularization (Figure 1B).

**Figure 1.**
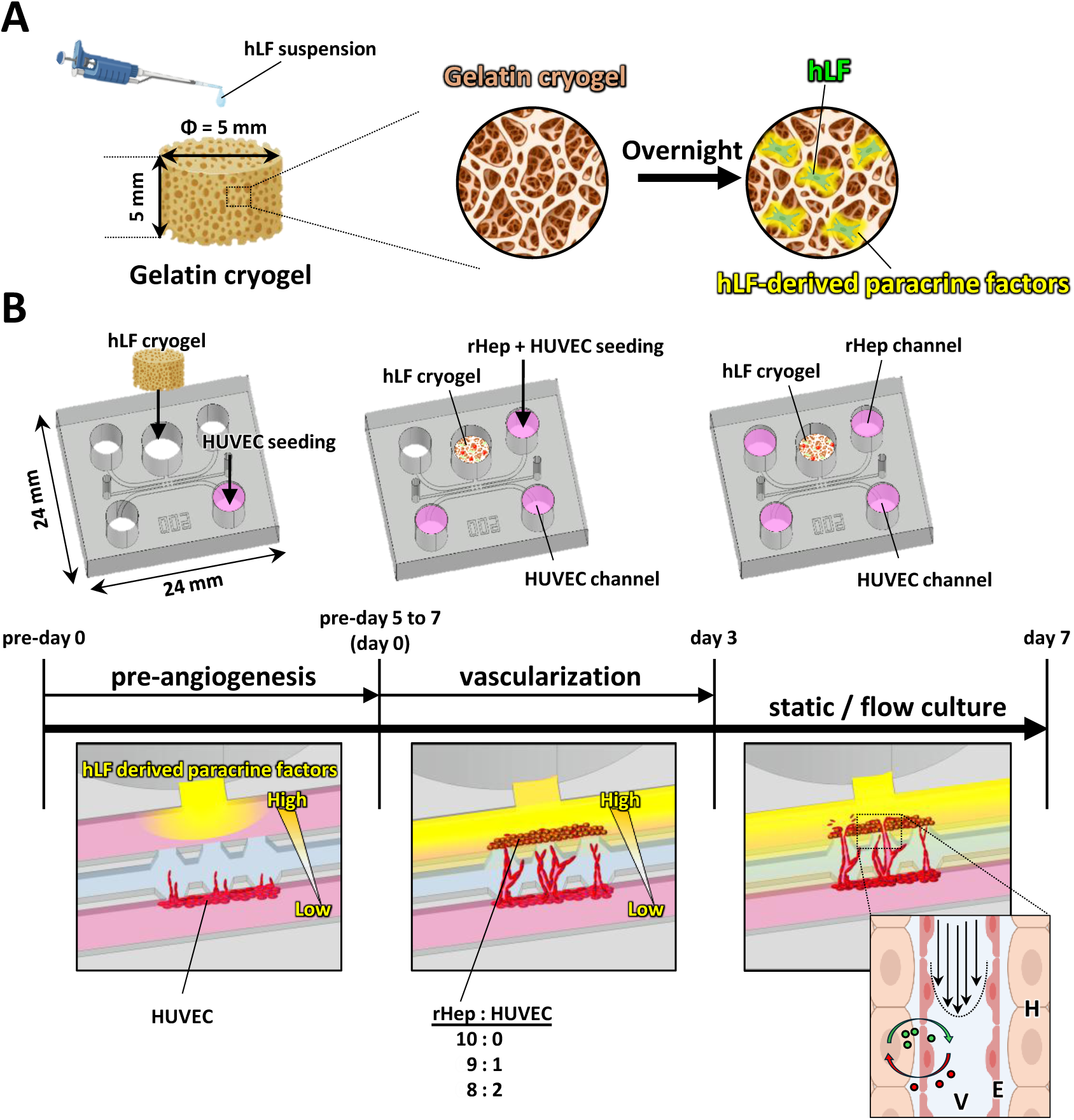
Schematic illustrations of the paracrine factor local gradient (PFLG) generating system for constructing vascularized hepatocyte tissue. A: Preparation of hLF-loaded gelatin cryogel (hLF cryogel). B: Experimental timeline for the construction of vascularized hepatocyte tissue. The PFLG generated by the hLF cryogel induces angiogenesis from the HUVEC channel (pre-angiogenesis from pre-day 0 to pre-day 5–7). Rat primary hepatocytes (rHeps) and HUVECs are then seeded into the rHep channel on day 0. From day 0 to day 3, microvessels gradually extend toward the hepatocyte tissue and eventually penetrate it. From day 3 to day 7, cells are cultured under either static or flow conditions. The magnified schematic illustration in the right panel shows the architecture of vascularized hepatocyte tissue (H: hepatocytes, E: endothelial cells, V: microvessel).

First, we characterized hLF culture within the gelatin cryogels. hLFs were seeded onto gelatin cryogels, which could be easily handled using forceps (**Figure 2A**). The physicochemical properties of the cryogels influence the efficiency of paracrine factor delivery. The internal structure of the cryogels was visualized via autofluorescence imaging using a confocal microscope. Cross-sectional images revealed an interconnected porous architecture with uniform pore distribution and an average pore size of 62.4 ± 17.4 µm (Figure 2B). The cryogels also exhibited a porosity of 62.8 ± 1.9% and a swelling ratio of 44.3 ± 2.5%. These properties support efficient nutrient transport and paracrine factor diffusion while providing sufficient space for hLF growth and proliferation.^[29,30]^

**Figure 2.**
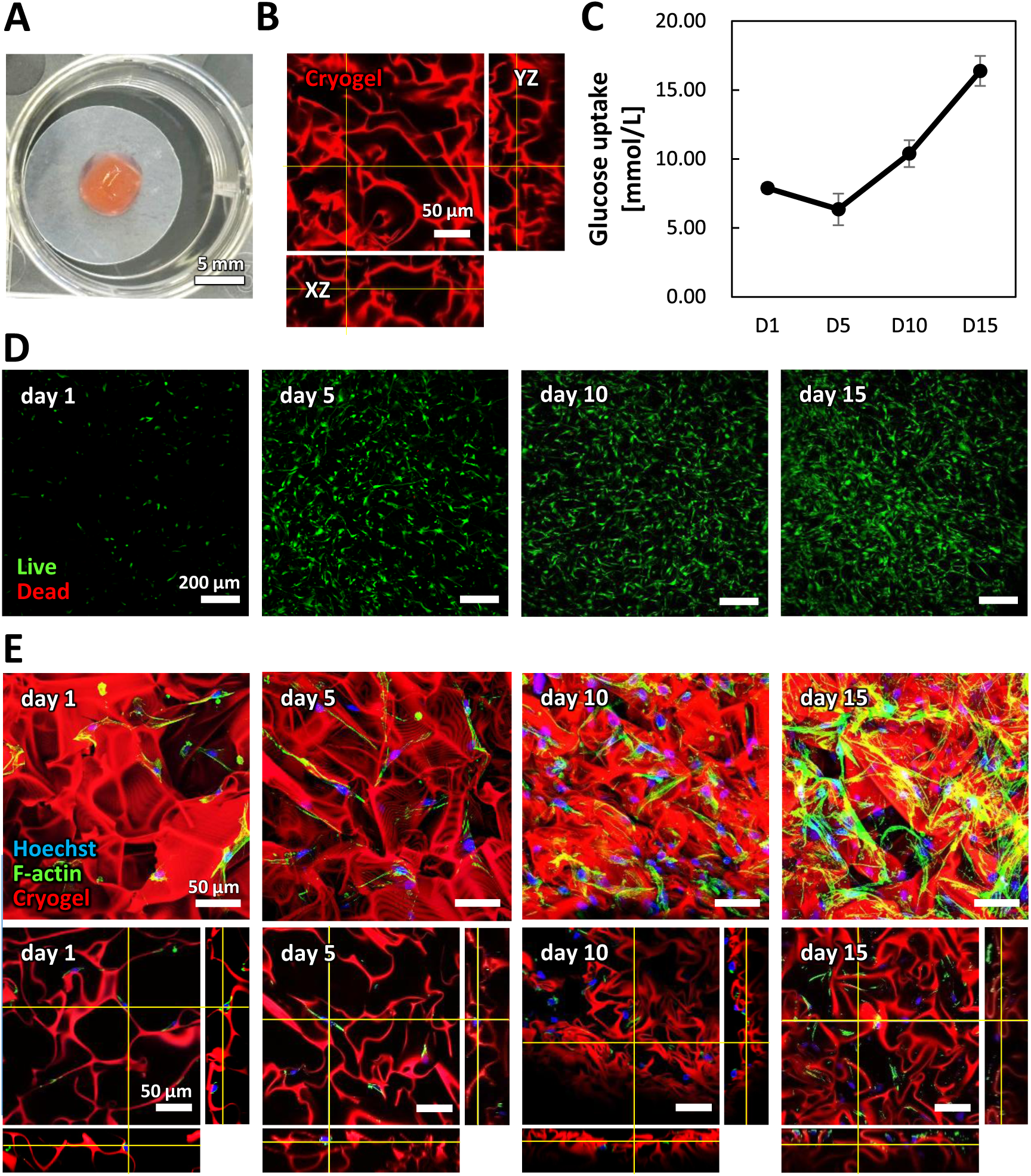
Characterization of hLF culture in gelatin cryogels. A: Photograph of an hLF cryogel. Scale bar, 5 mm. B: Confocal images showing the porous structure of the gelatin cryogel (red: cryogel autofluorescence). Scale bar, 50 µm. C: Quantitative analysis of glucose uptake by hLFs in cryogels (10,000 cells cryogel^−1^) over 15 days. D: Projection images of hLF live/dead staining in cryogels during days 1–15 (green: calcein-positive live cells; red: EthD-III-positive dead cells). Scale bars, 200 µm. E: Immunofluorescence images of hLF cryogels from day 1 to day 15. Blue: cell nuclei (Hoechst 33342), green: F-actin (Alexa Fluor 488-phalloidin), red: cryogel autofluorescence. Upper panels show projection images, and lower panels show cross-sectional images of hLF cryogels. Scale bars, 50 µm.

Next, we evaluated hLF cultivation within the cryogels to confirm their suitability as cell carriers. hLF growth kinetics were monitored by glucose uptake analysis over a 15-day culture period (Figure 2C). Glucose uptake was initially elevated on day 1, possibly reflecting the energy demand associated with cell attachment and adaptation to the cryogel environment. Although glucose uptake slightly decreased by day 5, it subsequently increased progressively until day 15, indicating sustained cell proliferation and metabolic activity within the cryogel matrix (Figure 2C).

Live/dead staining was also performed to assess cell viability. Fluorescence images revealed a progressive increase in hLF density, with the majority of cells remaining viable and minimal cell death observed at all time points (Figure 2D). These results were consistent with the glucose uptake profile, confirming that the elevated glucose consumption on day 1 was attributable to the energetic demands of cell attachment rather than cell death.

To further investigate cell morphology and organization, actin filaments and nuclei of hLFs cultured within cryogels were stained (Figure 2E). Projection images showed that actin filaments in hLFs gradually extended over time. Cross-sectional images on day 1 confirmed that the cryogels provided a supportive microenvironment for hLF adhesion. Furthermore, sequential cross-sectional images from day 1 to day 15 revealed progressive cryogel deformation corresponding to hLF extension over time.

### Angiogenic response optimization based on initial hLF cell density within the cryogels

The angiogenic response was regulated by the magnitude of the PFLG derived from hLFs, which can be enhanced by increasing the number of hLFs within the cryogels.^[28]^ In our PFLG-based culture system, HUVECs were initially cultured alone to establish microvascular networks prior to coculture with hepatocytes for vascularization. To investigate the effect of hLF density in the cryogels during this pre-angiogenesis phase, we prepared cryogels with different initial hLF densities and evaluated their proangiogenic performance on day 5 (referred to as Pre-day 5) (**Figure 3A**, Figure S1 Pre-angiogenesis phase).

**Figure 3.**
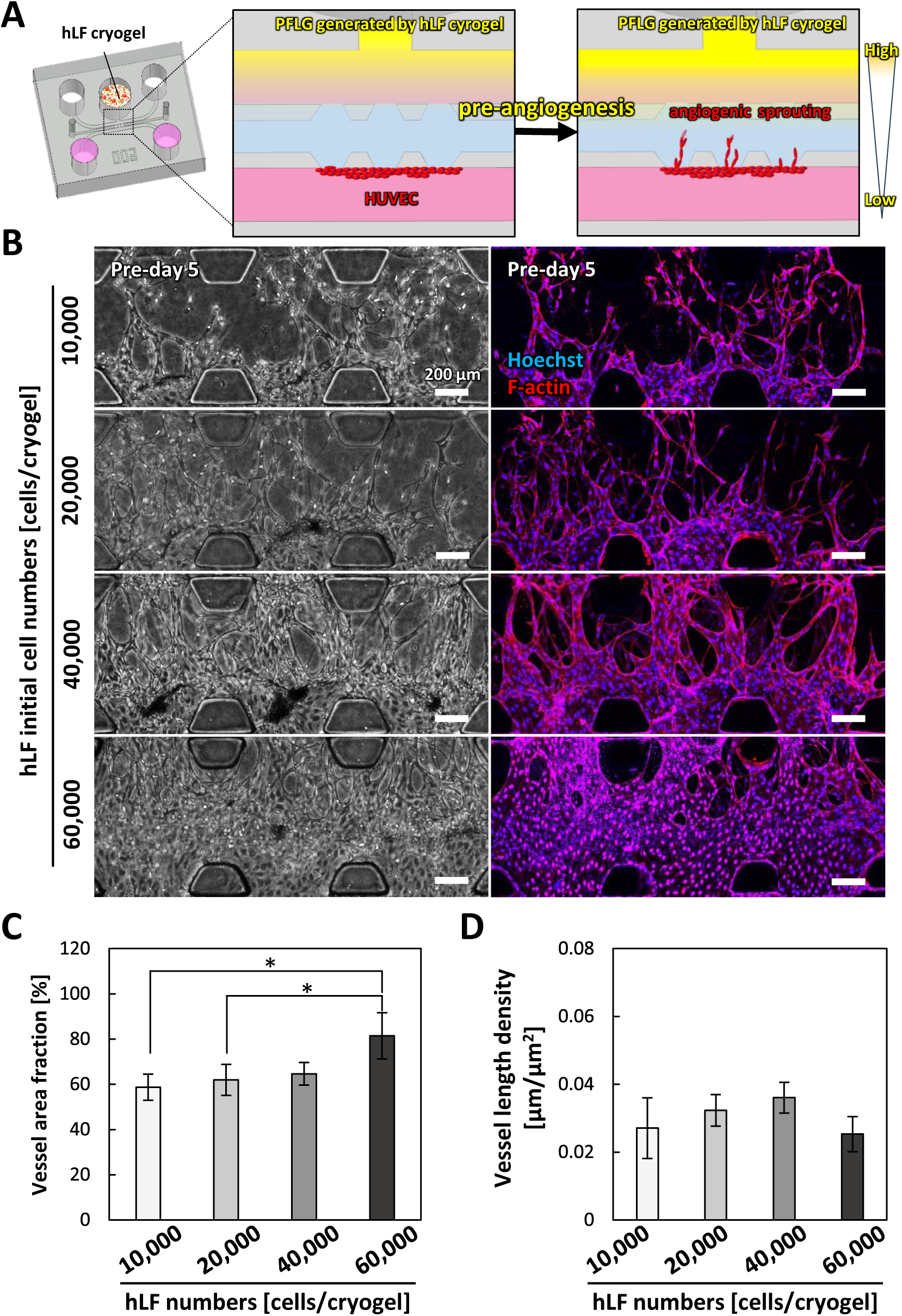
Angiogenic responses depending on the initial hLF number in cryogels. A: Schematic illustrations of PFLG generated by hLF cryogels. Angiogenic sprouting was induced by the PFLG (pre-angiogenesis). B: Representative phase-contrast and fluorescent projection images of microvessels on pre-day 5 with different initial hLF numbers seeded in the cryogel. Blue: cell nuclei (Hoechst 33342), red: F-actin (Alexa Fluor 594-phalloidin). Scale bars, 200 μm. C: Quantitative analysis of vessel area on pre-day 5. D: Quantitative analysis of vessel density on pre-day 5. Data are presented as mean ± SD. \**P* < 0.05, \*\**P* < 0.01, \*\*\**P* < 0.001 (One-way ANOVA followed by Tukey’s HSD test).

Phase-contrast and fluorescence imaging revealed a dose-dependent angiogenic response corresponding to increasing hLF densities from 10,000 to 60,000 cells cryogel^−1^. However, at the highest density (60,000 cells cryogel^−1^), microvessel structures were not maintained due to the fusion of adjacent vessels (Figure 3B). Quantitative analysis confirmed that the vessel area fraction increased with higher hLF densities (Figure 3C). In contrast, vessel length density peaked at 40,000 cells cryogel^−1^, while the 60,000 cells cryogel^−1^ condition exhibited reduced vessel length density, likely due to microvessel fusion (Figure 3D). Based on these findings, a density of 40,000 cells cryogel^−1^ was selected for subsequent experiments involving vascularization of hepatocyte tissue.

### Vascularization of hepatocyte tissue depending on rHep-to-HUVEC ratios

Following the pre-angiogenesis phase, rHeps were seeded into the rHep channel to construct vascularized hepatocyte tissue (**Figure 4A**, Figure S1 Vascularization phase). To optimize vascularization, three different rHep-to-HUVEC ratios (10:0, 9:1, and 8:2) were evaluated. Projection images on day 5 revealed microvessels penetrating into the hepatocyte tissue. Many penetrating microvessels were observed at the 9:1 rHep-to-HUVEC ratio (Figure 4B, arrowheads), whereas the 10:0 condition showed almost no penetrating microvessels. In this condition, unpenetrated microvessels displayed fragmented and discontinuous structures, which are signs of regression. The enlarged images also revealed dead cells within the hepatocyte tissue (Figure 4B, asterisks). The 8:2 condition resulted in a limited number of penetrating microvessels (Figure 4B, arrowheads). Notably, a higher proportion of HUVECs did not further enhance vessel penetration but rather led to the formation of isolated endothelial clusters (Figure 4B).

**Figure 4.**
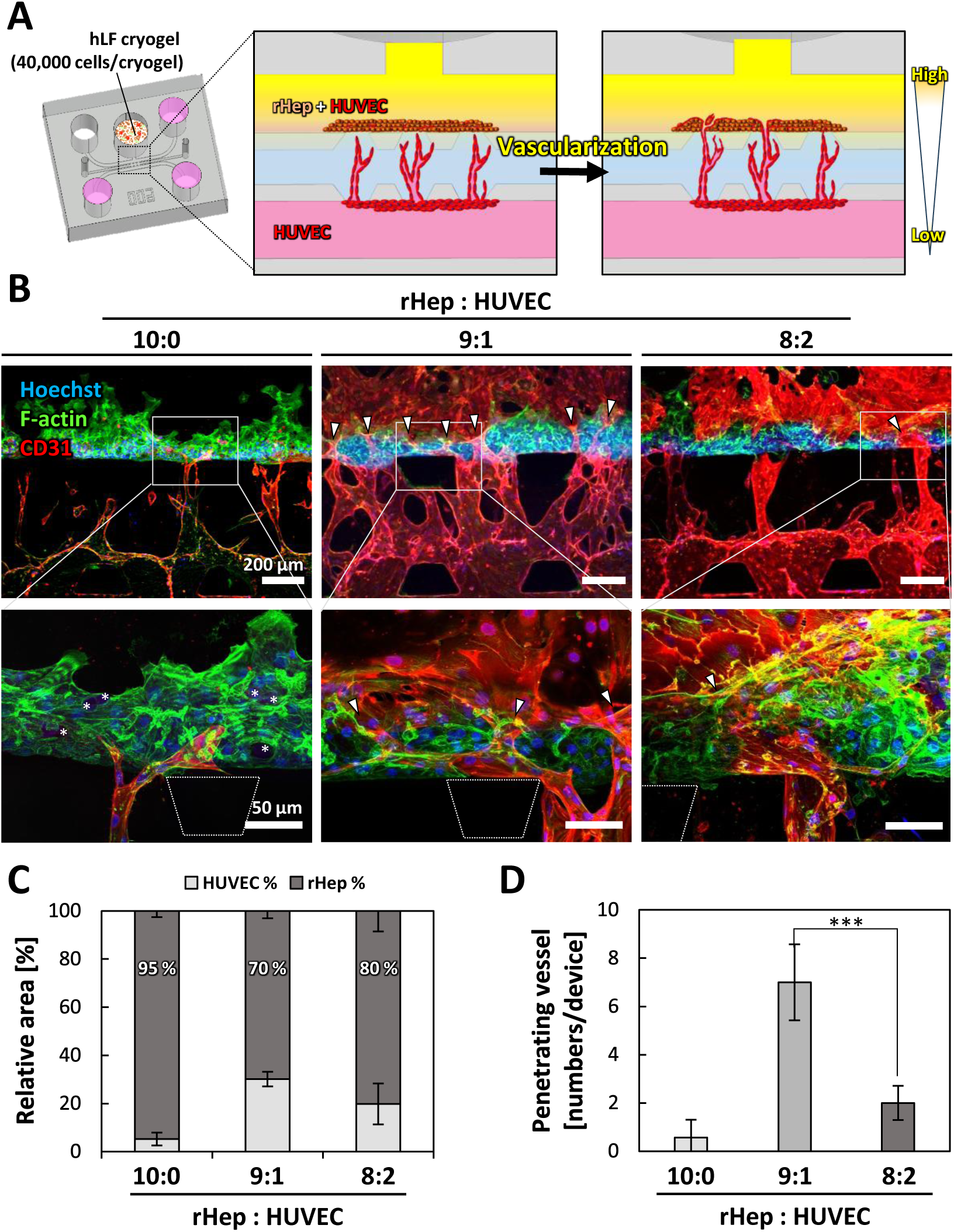
Vascularization of hepatocyte tissue depending on the rHep:HUVEC ratio. A: Schematic illustrations of PFLG generated by hLF cryogel, inducing microvessel penetration into hepatocyte tissue. B: Fluorescent projection images of vascularized hepatocyte tissue on day 5. Blue: cell nuclei (Hoechst 33342), green: F-actin (Alexa Fluor 488-phalloidin), red: HUVECs (CD31). Arrowheads indicate microvessels penetrating hepatocyte tissue. Asterisks in the magnified image indicate dead cells in hepatocyte tissue. Scale bars, 200 µm (upper panel); 50 µm (lower panel, magnified images). C: Quantitative analysis of vessel area ratio in fluorescent projection images of vascularized hepatocyte tissues on day 5. D: Quantitative analysis of the number of penetrating vessels on day 5. Data are presented as mean ± SD. \**P* < 0.05, \*\**P* < 0.01, \*\*\**P* < 0.001 (One-way ANOVA followed by Tukey’s HSD test).

Quantitative analysis revealed hepatocyte coverage of 95%, 70%, and 80% in the 10:0, 9:1, and 8:2 rHep-to-HUVEC ratio conditions, respectively (Figure 4C). Among the tested conditions, the 9:1 condition exhibited the highest number of penetrating microvessels, which was significantly greater than that observed in the 8:2 condition (Figure 4D). Based on these findings, the 9:1 condition was selected for subsequent experiments.

### Vascularized hepatocyte tissue recapitulating hepatic sinusoid-like architecture

The architecture of the vascularized hepatocyte tissue was examined by immunofluorescence staining. Projection images revealed that the tissue was penetrated by microvessels (**Figure 5A**). Cross-sectional images confirmed that microvessels traversed the entire thickness of the hepatocyte tissue (Figure 5B, arrowheads). The YZ cross-sectional view clearly showed a continuous microvessel penetrating the tissue (Figure 5B, arrowheads), and a magnified XZ view demonstrated that the microvessel was surrounded by rHeps in close proximity, forming sinusoid-like structures (Figure 5B, asterisks). Notably, extensive bilayer F-actin structures were observed between rHeps in the vascularized tissue, suggesting potential restoration of hepatic polarity upon vascularization.^[31]^ 3D reconstruction images and a corresponding video were also generated to support these observations (Figure S2, Video S1, Supporting Information). This architecture closely resembled native hepatic sinusoids, in which hepatocytes maintain direct contact with SECs (Figure 5C).

**Figure 5.**
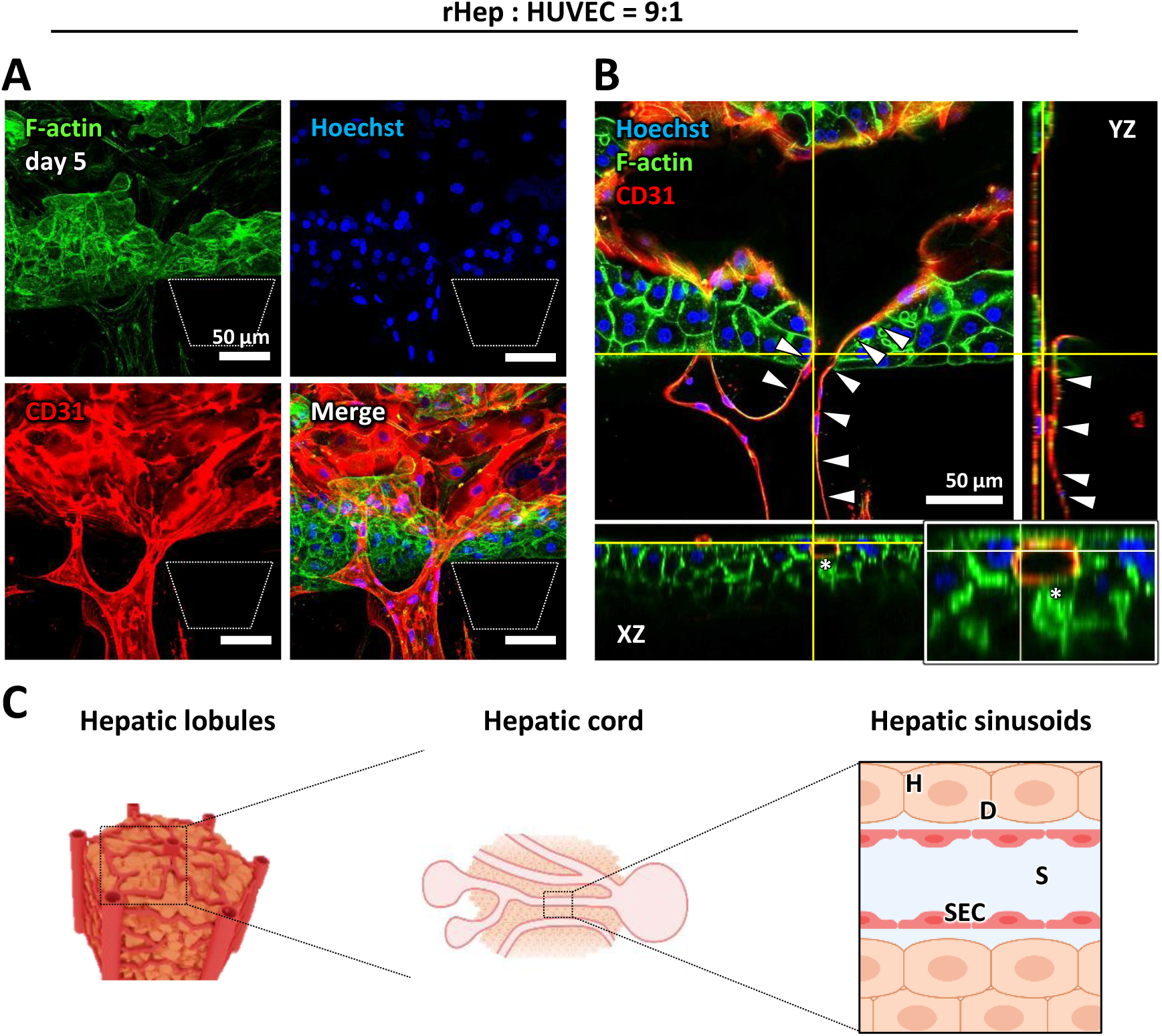
Morphological analysis of vascularized hepatocyte tissue. A: Immunofluorescence confocal images of vascularized hepatocyte tissue (rHep:HUVEC = 9:1) on day 5. Blue: cell nuclei (Hoechst 33342), green: F-actin (Alexa Fluor 488-phalloidin), red: HUVECs (CD31). B: Cross-sectional images of vascularized hepatocyte tissue. Arrowheads indicate microvessels penetrating the hepatocyte tissue. The asterisk in the magnified XZ section highlights hepatic sinusoid-like structures, where hepatocytes are in close contact with microvessels. Scale bar, 50 µm. C: Schematic illustrations of hepatic sinusoids (H: hepatocytes, D: space of Disse, SEC: sinusoid endothelial cells, S: hepatic sinusoid).

### Hepatocyte polarity and function in vascularized hepatocyte tissue

As we hypothesized that vascularization would enhance hepatocyte polarity and function, we evaluated both parameters under different culture conditions. Immunofluorescence staining was performed to assess hepatocyte polarity, with particular focus on MRP2, a marker of the bile canalicular domain. As control conditions, hepatocytes were cultured without hLF-loaded cryogel (hLF (−)) and without the pre-angiogenesis phase (Pre-angio (−)). Under these conditions, MRP2 expression was not detectable in rHeps (**Figure 6A**). However, the addition of hLF-loaded cryogel (hLF (+), Pre-angio (−)) resulted in weak and sparse MRP2 expression between rHeps (Figure 6A, arrowheads). Notably, with the inclusion of the pre-angiogenesis phase (hLF (+), Pre-angio (+)), abundant MRP2 expression was observed between rHeps within the vascularized hepatocyte tissue. The MRP2-positive bile canaliculi were aligned in parallel with the penetrating microvessels (Figure 6A). Quantitative analysis revealed a significantly higher ratio of MRP2-positive area in the vascularized hepatocyte tissue compared to the other two conditions (Figure 6D). These findings clearly demonstrate enhanced rHep polarization.

**Figure 6.**
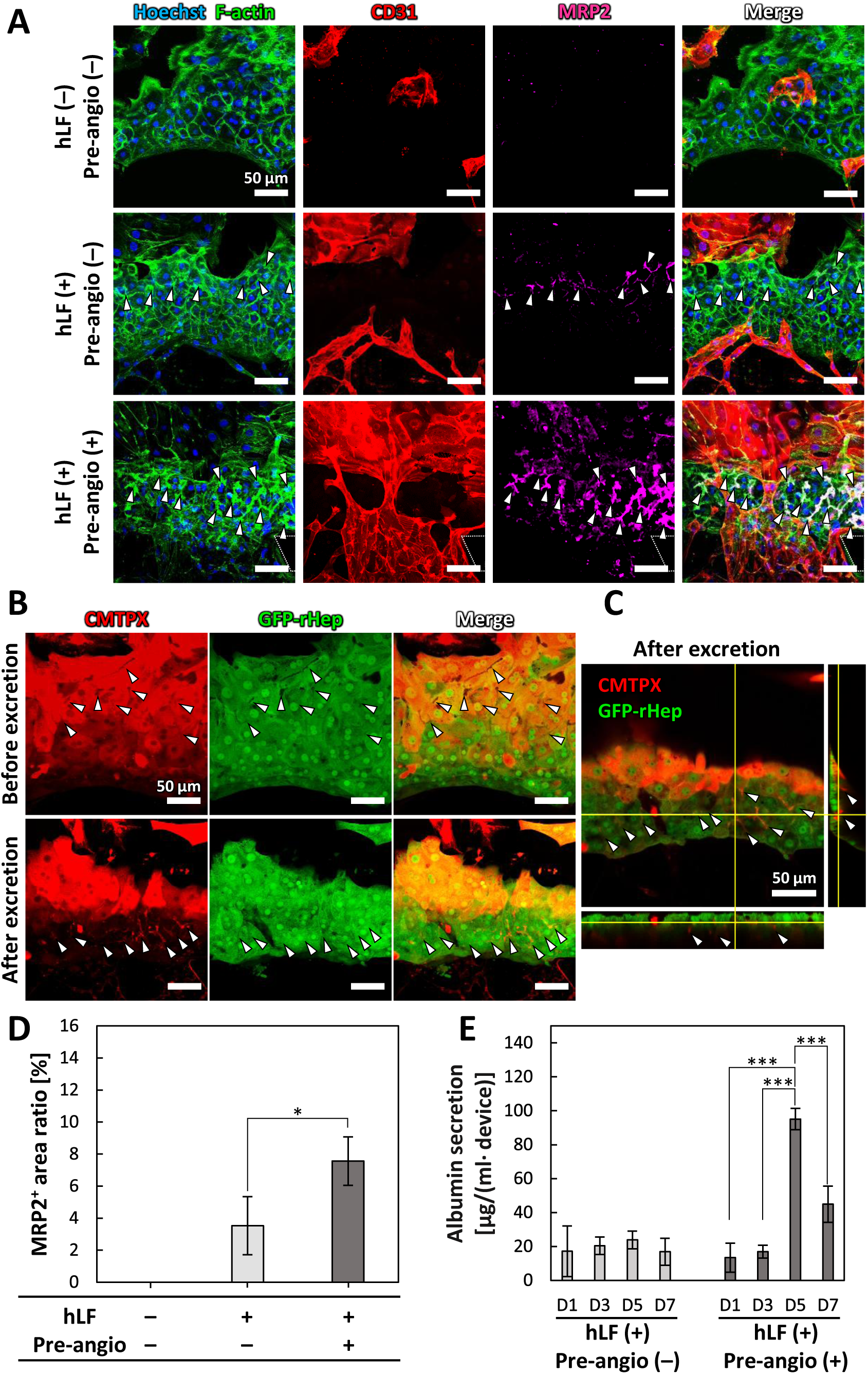
Assessment of hepatic functions under static culture. A: Immunofluorescence projection images of bile canalicular formation in vascularized hepatocyte tissue. Projection images were constructed from Z-stack confocal images. Blue: cell nuclei (Hoechst 33342), green: F-actin (Alexa Fluor 488-phalloidin), red: HUVECs (CD31), pink: bile canaliculi (MRP2). Arrowheads indicate MRP2-positive bile canaliculi. B: Excretion of CellTracker Red CMTPX by polarized hepatocytes in vascularized hepatocyte tissue (red: CMTPX; green: GFP-rHep). Arrowheads indicate CMTPX excreted into bile canaliculi. C: Cross-sectional image showing CMTPX excretion into bile canaliculi in vascularized hepatocyte tissue (red: CMTPX; green: GFP-rHep). Arrowheads indicate CMTPX in bile canaliculi. Scale bars, 50 µm. D: Quantitative analysis of MRP2-positive area ratio in hepatocyte tissue. E: Quantification of albumin secretion from hepatocyte tissues. Data are presented as mean ± SD. \**P* < 0.05, \*\**P* < 0.01, \*\*\**P* < 0.001 (One-way ANOVA followed by Tukey’s HSD test).

To further investigate the function of bile canaliculi in the vascularized hepatocyte tissue, we added CellTracker Red CMTPX to the culture medium. This dye is spontaneously taken up by rHeps, hydrolyzed to a fluorescently active form in the cytoplasm, and subsequently excreted into bile canaliculi.^[19]^ Projection images demonstrated that CMTPX was first internalized by rHeps and accumulated in the cytoplasm (Figure 6B, CMTPX). Meanwhile, bile canaliculi appeared as distinct black line structures (Figure 6B, arrowheads) between rHeps prior to excretion.^[19]^ After 10 min, CMTPX was excreted from the cytoplasm into bile canaliculi (Figure 6B, arrowheads), thereby visualizing their structure (Figure 6B, Video S2, Supporting Information). Cross-sectional images further revealed the 3D structure of the excreted CMTPX within the bile canaliculi (Figure 6C, arrowheads).

Next, we evaluated hepatocyte function by measuring albumin secretion from rHeps using an enzyme-linked immunosorbent assay (ELISA) over a 7-day culture period. Under vascularized conditions (hLF (+), Pre-angio (+)), albumin secretion levels markedly increased by day 5, reaching 7.1-fold higher than those on day 1 (Figure 6E). However, by day 7, the levels slightly declined to 3.3-fold above day 1. In contrast, hepatocyte tissue cultured without the pre-angiogenesis phase (hLF (+)/pre-angio (−)) exhibited consistently low albumin secretion throughout the culture period, highlighting the critical role of vascularization in supporting hepatic function.

### Functional perfusion analysis of vascularized hepatocyte tissue

In this study, we developed the PFLG-generating system to construct perfusable hepatocyte tissue. Although the results described above demonstrated the successful construction of vascularized hepatocyte tissue, the culture was conducted under static conditions. Therefore, we further investigated whether the vascularized hepatocyte tissue supports functional perfusion. To this end, fluorescently labeled microbeads (diameter: 1 µm) were introduced into the culture medium to visualize flow through the penetrating microvessels. Time-lapse imaging captured the real-time movement of the microbeads within the microvessels, confirming successful perfusion through the vascularized hepatocyte tissue (**Figure 7A**, Videos S3 and S4, Supporting Information).

**Figure 7.**
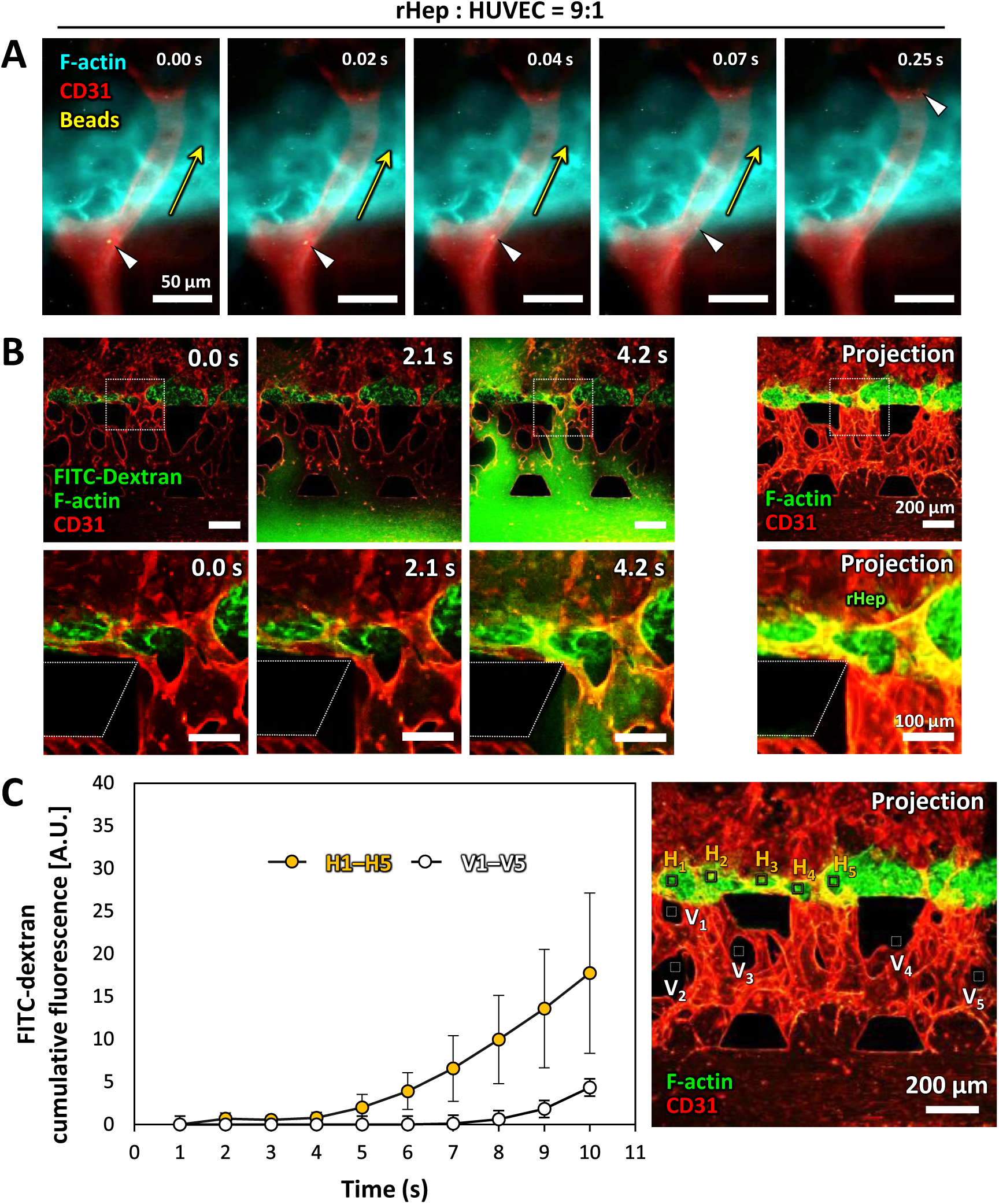
Functional perfusion analysis of vascularized hepatocyte tissue on day 5 of the vascularization phase. A: Time-lapse images of fluorescently labeled microbeads (diameter: 1 µm) perfused through vascularized hepatocyte tissue. Arrowheads indicate a perfused microbead, and arrows indicate the flow direction within the microvessel. Scale bar, 50 µm. B: Time-lapse and projection images showing perfusion of 70 kDa FITC-dextran solution through vascularized hepatocyte tissue. Green: FITC-dextran and F-actin (Alexa Fluor 488-phallidin), red: microvessels (CD31). Scale bars, 200 µm (upper panel); 100 µm (lower panel, magnified images). C: Quantitative analysis of cumulative FITC-dextran fluorescence in regions H1–H5 and V1–V5 shown in the right image. Boxes in the projection image indicate regions of interest for analyzing cumulative FITC-dextran fluorescence during the perfusion process (H1–H5: hepatocyte tissue regions; V1–V5: hydrogel regions)

To further assess the functional perfusion of the vascularized hepatocyte tissue, we performed comprehensive perfusion analyses using FITC-dextran solution (Figure 7B). The perfusion of FITC-dextran visualized molecular transport dynamics through the microvascular network. Time-lapse fluorescence imaging showed that the FITC-dextran solution was introduced into the HUVEC channel. The fluorescent signal propagated through the microvascular network within the hydrogel and eventually perfused the microvessels penetrating the 3D hepatocyte tissue formed in the rHep channel. The images showed FITC-dextran leakage from the microvessels, with greater accumulation within the hepatocyte tissue than in the surrounding hydrogel (Figure 7B, Video S2, Supporting Information). Quantitative analysis clearly demonstrated the fluorescence intensity differences between the hepatocyte tissue (Figure 7C, ROIs H1–H5) and hydrogel regions (Figure 7C, ROIs V1– V_5_). After 10 sec, the average intensity in the hepatocyte tissue region was approximately 3-fold higher than that in the hydrogel region, supporting our observation.

### Induction of active rHep proliferation in vitro under perfusion conditions

Finally, we investigated hepatocyte proliferation in vascularized hepatocyte tissues under static and perfusion culture conditions (**Figure 8A**). Phase-contrast imaging showed tissue growth during days 1–5, suggesting potential hepatocyte proliferation (Figure S3, Supporting Information). To verify this, we performed Ki67 staining to identify proliferating cells. Non-vascularized hepatocyte tissues (rHep:HUVEC ratio = 9:1) showed few Ki67-positive rHeps only when directly contacted by mature microvessels, while tissues without vascular contact remained quiescent (Figure S4, Supporting Information). Under static conditions of vascularized hepatocyte tissue, fluorescence images showed few Ki67-positive rHeps, with no significant difference regardless of the presence of hLF cryogel on days 5 and 7. In contrast, under perfusion culture, Ki67-positive rHeps markedly increased, including both mononuclear and binuclear cells (Figure 8B, arrowheads). Magnified images revealed double-layered F-actin structures between proliferating rHeps (Figure 8B, asterisks). Notably, hepatocytes undergoing mitosis were observed under 1 mmH_2_O perfusion on day 7 (Figure 8C).

**Figure 8.**
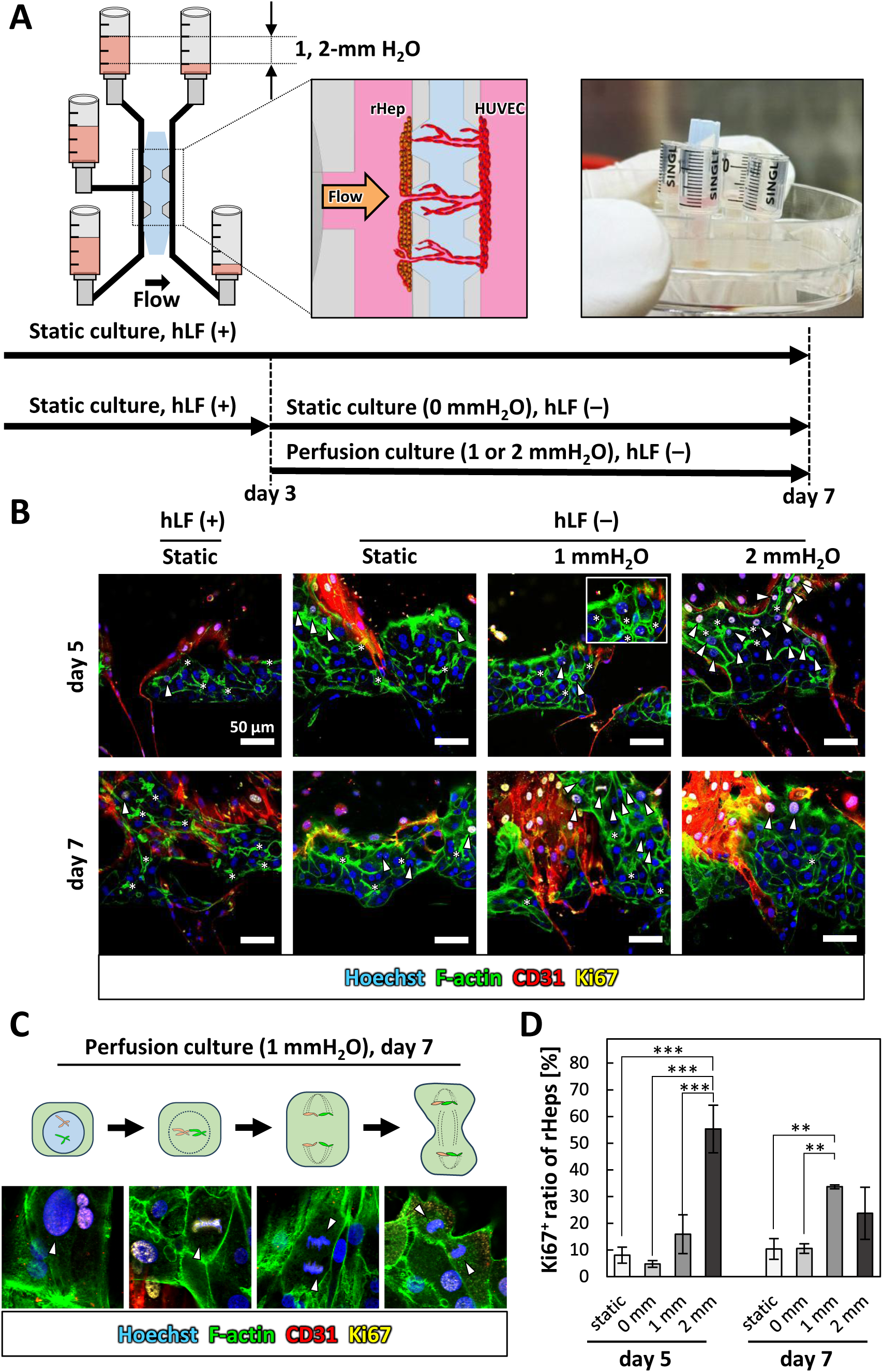
Hepatocyte proliferation responses in vascularized hepatocyte tissue under perfusion culture. A: Schematic illustrations of static and perfusion culture conditions. Cells were cultured with hLF cryogels (hLF(+)) until day 3. From day 3 to day 7, some cells were cultured without hLF cryogels (hLF(−)) under either static (0 mmH_2_O) or perfusion (1 or 2 mmH_2_O) conditions. B: Immunofluorescence projection images of vascularized hepatocyte tissue under static and perfusion culture. Projection images were constructed from Z-stack confocal images. Blue: cell nuclei (Hoechst 33342), green: F-actin (Alexa Fluor 488-phalloidin), red: HUVECs (CD31), yellow: proliferating cell nuclei (Ki67). Arrowheads indicate Ki67-positive proliferating hepatocytes. Asterisks indicate bile canaliculi-like structures, showing double-layered F-actin between hepatocytes. Scale bars, 50 µm. C: Representative immunofluorescence images showing Ki67-positive rHeps undergoing mitosis. Arrowheads indicate both interphase nuclei and mitotic chromosomal dynamics. Blue: cell nuclei (Hoechst 33342), green: F-actin (Alexa Fluor 488-phalloidin), red: HUVECs (CD31), yellow: proliferating cell nuclei (Ki67). D: Quantification of the ratio of Ki67-positive rHeps. Data are presented as mean ± SD. \**P* < 0.05, \*\**P* < 0.01, \*\*\**P* < 0.001 (One-way ANOVA followed by Tukey’s HSD test).

To quantify rHep proliferation, the ratio of Ki67-positive rHeps was measured. The results confirmed significantly higher Ki67-positive ratios under perfusion culture on both days 5 and 7. On day 5, the ratio under 1 mmH_2_O perfusion was 2.0-fold higher than that under static culture with hLF-loaded cryogels (Figure 8D, static), and 3.4-fold higher than that under static culture without hLF-loaded cryogels (Figure 8D, 0 mm). Furthermore, the Ki67-positive ratio under 2 mmH_2_O perfusion culture was significantly higher, reaching 3.6-fold that of 1 mmH_2_O perfusion. On day 7, the Ki67-positive ratio under 1 mmH_2_O perfusion increased further, reaching approximately 3.4-fold higher than static culture. However, under 2 mmH_2_O perfusion, the Ki67-positive ratio slightly decreased on day 7 (Figure 8D).

## Discussion

### PFLG-generating system for pre-angiogenesis and vascularization

In this study, we developed a PFLG-generating system that promotes both pre-angiogenesis and vascularization of hepatocyte tissues. The system integrates hLF-loaded cryogels with a microfluidic platform, wherein the cryogels act as a sustained source of paracrine factors. Cryogels are highly porous biomaterials that serve as effective hLF carriers.^[29,30,32]^ This feature enables continuous delivery of paracrine factors, thereby establishing a PFLG within the device.

Conventional strategies typically add exogenous growth factors (e.g., VEGF) to the culture medium, providing only transient stimulation that decays over time.^[33–35]^ Moreover, maintaining a stable spatial concentration gradient of growth factors within the culture microenvironment is challenging. By contrast, direct coculture approaches introduce uncontrolled cell–cell contacts over time. Notably, excessive hLF proliferation in such systems has been associated with pro-fibrotic remodeling.^[36–38]^

The PFLG-generating system addresses these limitations by providing sustained paracrine stimulation that scales with hLF proliferation, consistent with our previous findings.^[28]^ At the same time, it maintains physical separation between cell populations (e.g., rHeps/HEVECs vs. hLFs) while creating a targeted gradient that guides microvessels to penetrate hepatocyte tissue. Importantly, the hLF-loaded cryogel can be physically removed, permitting on-demand switching of culture conditions according to the stage of tissue morphogenesis.

### PFLG-mediated direct microvessel penetration of 3D hepatocyte tissue

Fibroblasts secrete multiple pro-angiogenic factors (e.g., VEGF, FGF, and PDGF) that enhance vascular formation.^[39–41]^ These paracrine factor cocktails regulate endothelial cell proliferation, migration, and lumen formation via downstream signaling, specifically the MAPK/ERK and PLCγ/PKC pathways.^[42]^ The hLF-loaded cryogel continuously releases such factors, establishing a PFLG across the hydrogel from the rHep channel toward the HUVEC channel of the microfluidic device. This gradient promotes sprout formation^[34,39]^ and guides microvascular extension toward the rHep channel. The platform enables a pre-angiogenesis phase in which microvascular networks develop before hepatocyte seeding; consequently, during subsequent 3D tissue formation, microvessels penetrate the nascent hepatocyte tissue. Timely penetration is critical because rHeps establish strong cell–cell adhesions within approximately 3 days post-seeding, after which these adhesions impede further vascular extension (Fig. S4, hLF(+) Angio (+) Penetrated (–)), consistent with our previous report.^[43]^ This temporal constraint underscores the need to optimize angiogenic conditions. In our pre-angiogenesis experiments, the angiogenic response scaled with the initial hLF density, peaking at 40,000 hLFs cryogel^−1^. During the vascularization phase, a 9:1 rHep:HUVEC ratio most effectively supported microvessel penetration. These proportions closely mimic those of native liver tissue, where hepatocytes comprise approximately 70% and the sinusoidal space occupies 30% of the tissue area, ^[44]^ indicating that relatively few endothelial cells are sufficient for vascular guidance. Consistent with previous studies, the incorporation of endothelial cells enhanced the vascularization of hepatocyte tissues.^[45]^

Nevertheless, achieving intimate hepatocyte–microvessel contact remains challenging in many hepatic models. Needle-based templating is widely used but is constrained by needle diameter, limiting vascular scale.^[46]^ While bioprinting can mitigate channel-size constraints, hepatocytes and vessels are frequently separated by hydrogel matrices with gaps exceeding tens of micrometers.^[47]^ Our PFLG-generating system overcomes these barriers by driving direct microvessel penetration into hepatocyte tissue, thereby recapitulating sinusoid-like architecture in vitro.

### Vascularized hepatocyte tissue with sinusoid-like architecture enhances hepatic functions

Hepatic sinusoids possess a unique architecture characterized by direct hepatocyte–endothelial contact, enabling efficient substance exchange between blood and liver in vivo.^[48–50]^ As reported previously, we replicated sinusoid-like architecture in vitro by vascularizing rHep–hLF spheroids.^[51]^ Building on this foundation, the present study optimized hepatocyte–endothelial cell ratios and guided angiogenic directionality using the PFLG-generating system. Immunofluorescence revealed continuous microvessels penetrating hepatocyte tissue, and orthogonal cross-sections showed luminal microvessels closely apposed to rHeps. The reconstructed microarchitecture facilitates mass transport and rHep–HUVEC communication, which are critical for enhancing hepatic functions.^[2]^

Polarity is a hallmark of hepatocellular differentiation.^[52–54]^ Immunofluorescence for MRP2 showed apical localization at canalicular membranes between adjacent rHeps, indicating restoration of polarity and formation of bile canaliculi. Vascularized hepatocyte tissues exhibited abundant bile canaliculi radially aligned with microvessels, suggesting that rHeps recognized apical (canalicular) and sinusoidal domains. Consistent with MRP2 localization, albumin secretion increased significantly from day 5 onward, following microvessel penetration of the hepatocyte tissue. By contrast, non-vascularized tissues displayed limited bile canaliculus formation and significantly lower albumin levels, despite exposure to paracrine factors from the hLF-loaded cryogel and rHep-conditioned media. Together, these results indicate that vascularization enhances hepatocyte function through at least two mechanisms: (i) penetrating microvessels alleviate diffusion limitations by providing direct nutrient delivery and metabolic waste removal; and (ii) intimate rHep–microvessel apposition enables short-range exchange and delivery of endothelial-derived paracrine cues. These findings underscore the importance of vascularization for engineering functional hepatocyte tissues in vitro.

### Functional perfusion of vascularized hepatocyte tissues

Perfusion with 1-μm fluorescent microbeads and 70-kDa FITC–dextran demonstrated functional microvascular flow through microvessels traversing the vascularized hepatocyte tissue. Dextran leakage within hepatocyte-tissue regions was significantly greater than that in hydrogel regions, whereas microbeads exhibited no extravasation in either region. These findings indicate size-selective vascular permeability that permits macromolecular transport while excluding micrometer-scale particulates, consistent with the submicrometer fenestrations (approximately 50–400 nm) characteristic of SECs in vivo.^[48,55]^

Previous studies have demonstrated that HUVECs exhibit phenotypic plasticity in liver-mimetic microenvironments. Shaheen et al.^[56]^ reported that HUVECs cultured within decellularized liver scaffolds upregulated SEC markers and formed fenestrae-like structures, indicating a shift toward an SEC-like phenotype in response to liver ECM cues. De Haan et al.^[57]^ further showed that HUVECs retain transcriptional programs responsive to liver-specific signals, with partial induction of SEC markers upon transcription-factor overexpression. Together, these findings support the possibility that our vascularized hepatocyte tissue recapitulates key features of the sinusoidal microenvironment in vitro.

### Perfusion-triggered proliferation of hepatocytes in vascularized hepatocyte tissue in vitro

Hepatocyte behavior in vitro contrasts sharply with in vivo regeneration, in which proliferation typically accompanies dedifferentiation.^[58,59]^ Although 3D culture can preserve hepatocyte differentiation in vitro, proliferative capacity is typically lost.^[58]^ We therefore hypothesized that reconstructing the hepatic sinusoidal microenvironment in vitro would unlock rHep proliferation within 3D vascularized tissue.

Even under static conditions, the fraction of proliferating rHeps (Ki67-positive) reached approximately 10% in vascularized hepatocyte tissues. Two non-exclusive mechanisms may explain this observation. First, penetrating microvessels likely alleviate mass-transfer limitations, improving oxygen and nutrient delivery to rHeps. Second, daily medium exchange probably induces transient perfusion through the microvascular network, imposing endothelial shear that elicits the secretion of pro-mitogenic angiocrine factors.^[12,14,60]^ Consistent with this interpretation, flow-mediated mechanical cues have been reported to enhance hepatocyte cell-cycle entry.^[13,15]^

In contrast to static conditions, perfusion culture markedly enhanced hepatocyte proliferation in our vascularized hepatocyte tissues. Both mononucleated and binucleated rHeps exhibited cell-cycle entry, with the Ki67 labeling index increasing to 40–50%. Notably, rHeps in mitosis were directly observed under 1 mmH_2_O perfusion, suggesting the critical role of perfusion-mediated shear stress in activating hepatocyte proliferation. Perfusion-induced proliferation is plausibly mediated by endothelial mechanotransduction: Lorenz et al.^[12]^ showed that flow activates endothelial β1 integrin and VEGFR3, triggering SECs to release pro-mitogenic angiocrine factors (e.g., HGF, IL-6). Such mechanosensing provides a mechanistic framework for the perfusion-triggered hepatocyte proliferation observed in our PFLG-generating system.

The most significant achievement of this study is the induction of robust rHep proliferation while retaining differentiation and polarity within perfused hepatocyte tissue in vitro. Ki67-positive rHeps displayed a bilayer F-actin architecture between adjacent rHeps, which suggests the formation of bile canaliculi,^[61]^ under both perfusion and static culture conditions, indicating preserved bile canaliculi during proliferation. These in vitro observations recapitulate the in vivo phenomenon whereby hepatocytes maintain bile canaliculi while dividing.^[62]^

By contrast, prior organoid systems have relied on inflammatory cytokines (e.g., TNF-α, IL-6) to initiate hepatocyte proliferation in vitro,^[63,64]^ but typically require continuous cytokine exposure and show incomplete restoration of polarity. Moreover, the absence of functional vasculature constrains convective nutrient and oxygen transport, limiting sustained tissue growth. Chhabra et al. developed a platform integrating endothelial–hepatocyte interactions, in which perfusion combined with IL-1β stimulation modulated PGE_2_ signaling to initiate proliferation of cryopreserved hepatocytes.^[15]^ While Chhabra et al. demonstrated perfusion-induced hepatocyte proliferation via endothelial–hepatocyte interactions, their microfluidic system did not incorporate perfusable microvessels penetrating hepatocyte tissue. In contrast, our PFLG-generating platform enables direct parenchymal perfusion, allowing us to observe mitotic rHeps and assess nuclear localization and polarity during regeneration. In this study, we engineered a PFLG-generating platform that produces vascularized hepatocyte tissue perfused by microvessels penetrating the parenchyma. Importantly, we directly observed rHeps undergoing mitosis within these constructs, indicating that regenerative capacity can be engaged while preserving differentiation and polarity.

## Conclusion

In summary, we developed a PFLG-generating system that delivers sustained, spatially directed, and tunable paracrine stimulation during culture. Furthermore, the system provides programmable, multimodal control that transitions from paracrine-factor-driven angiogenesis to perfusion-mediated mechanotransduction. Its modular design enables real-time adjustment of cellular composition, paracrine profiles, stimulus intensity, and mechanical cues during culture. The resulting vascularized hepatocyte tissues formed functional microvessels resembling sinusoidal architecture while maintaining hepatic functions and proliferative capacity, thereby laying the foundation for scale-up toward transplantable liver tissues.

Despite these advances, several limitations of the platform require further optimization. First, hepatocytes must be protected from perfusion-induced shear stress; in our system, direct exposure of some rHeps to flow during perfusion culture led to deterioration in viability and function. Second, because we used freshly isolated primary rat hepatocytes, translation to human hepatocyte systems is a critical next milestone to enhance clinical relevance. Finally, a deeper mechanistic understanding of perfusion-mediated hepatocyte proliferation in vascularized hepatocyte tissue is needed to identify the key drivers and actionable targets.

## Experimental Section

### Preparation and characterization of gelatin cryogels

Gelatin cryogels were prepared as previously described,^[29]^ with minor modifications to the molding procedure and polymer concentration. Briefly, a 4% (w/v) gelatin solution was mixed with a 2% glutaraldehyde solution (GA, cross-linking agent) at a 1:1 volume ratio, immediately injected into syringe molds (1 ml), and incubated at –30°C overnight to induce cryogelation. The resulting cryogels were sliced into discs (5 mm thickness, Fig. 1A) and sequentially washed with 70% ethanol and distilled water (each overnight) to remove unreacted glutaraldehyde and byproducts. Cryogel porosity and swelling ratio were characterized as previously described.^[29]^ Briefly, pore sizes were determined using confocal microscopy based on the intrinsic autofluorescence of the cryogel structure. Porosity was determined using the ethanol displacement method by immersing dried cryogels in 99.5% ethanol, measuring the weight gain, and calculating porosity (%) according to Equation (1),

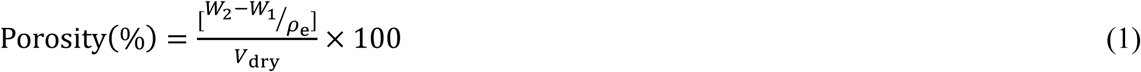

where *W*_1_ and *W*_2_ are the weights of dry and ethanol-immersed cryogels, *ρ*_e_ is the density of ethanol, and *V*_dry_ is the volume of the dry cryogel.

The swelling ratio was evaluated by immersing cryogels in distilled water at room temperature for 24 h to obtain the swollen weight (*W*_wet_), followed by freeze-drying to obtain the dry weight (*W*_dry_), and was calculated according to Equation (2).

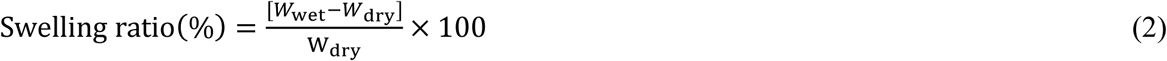

### Cell culture

Primary rat hepatocytes (rHeps) were isolated from Sprague-Dawley rats and green fluorescent protein (GFP) transgenic rats (250–350 g; Sankyo Labo Service, Tokyo, Japan) using a two-step collagenase perfusion method.^[19,65]^ Collagenase-digested liver cells were suspended in L-15 medium supplemented with 20 mM HEPES, 1.1 g l^−1^ galactose, 30 mg l^−1^ L-proline, 0.5 mg l^−1^ insulin, 10^−7^ M dexamethasone, and antibiotics. After centrifugation at 50*×g* for 1 min, performed twice, the pellet was resuspended in the L-15 medium and mixed with an equal volume of Percoll solution, consisting of 9 parts Percoll (Sigma-Aldrich, St. Louis, MO, USA) and 1 part 10× Hanks’ balanced salt solution without calcium and magnesium. The suspension was centrifuged at 50*×g* for 15 min to remove dead cells and nonparenchymal cells. The pellet was then resuspended in the medium and centrifuged again at 50*×g* for 1 min. Finally, the pellet was resuspended in Dulbecco’s Modified Eagle’s Medium (DMEM; Sigma-Aldrich, St. Louis, MO, USA) supplemented with 20 mM HEPES, 25 mM NaHCO_3_, 30 mg l^−1^ L-proline, 10% FBS, 0.5 mg ml^−1^ insulin, 10^−7^ M dexamethasone, 10 mM nicotinamide, 1 mM ascorbic acid 2-phosphate, 10 ng ml^−1^ epidermal growth factor (EGF), and antibiotics. To prevent degradation of the fibrin gel by plasmin secreted by rHeps, 6-aminocaproic acid (LKT Laboratories, USA) was added to the hepatocyte medium at 80 mg ml^−1^. Cell viability >90% was verified by the trypan blue exclusion test. All animal experiments were performed according to the Institutional Guidelines on Animal Experimentation at Keio University and approved by the Keio University Institutional Animal Care and Use Committee.

Human lung fibroblasts (hLFs; CC-2512, Lonza, MD, USA) and human umbilical vein endothelial cells (HUVECs; C2519A, Lonza, MD, USA) were commercially obtained. hLFs were cultured in fibroblast medium consisting of DMEM supplemented with 20 mM HEPES, 25 mM NaHCO_3_, 30 mg l^−1^ L-proline, 10% FBS, and 0.5 mg ml^−1^ insulin. HUVECs were maintained in endothelial growth medium (EGM-2; Lonza, MD, USA). Both cell types were expanded in collagen-coated culture dishes and used for subsequent experiments at passage 6.

Cell culture during the pre-angiogenesis phase (hLF-loaded cryogel-HUVEC coculture) was performed in a 1:1 mixture of the hepatocyte medium and EGM-2.^[43]^ For the vascularization phase (hLF-loaded cryogel-rHep-HUVEC triculture), a 1:1 mixture of the hepatocyte-conditioned medium (HepCM) and EGM-2 was used. All cultures were maintained in a humidified incubator at 37°C with 5% CO_2_. HepCM was employed because a previous study demonstrated its ability to enhance hepatocyte function.^[66]^ Consistent with this report, improved hepatocyte morphology was observed in 2D cultures supplemented with HepCM (Figure S5, Supporting Information). HepCM was prepared by culturing 2×10^6^ rHeps in 10 ml hepatocyte medium for 48 h. The conditioned medium was harvested by centrifugation at 2,600*×g* for 20 min, filtered through a 0.2 μm membrane to remove cellular debris, and stored at –30°C until use.

### hLF culture in cryogels

Cryogels were soaked in distilled water and sterilized by autoclaving. The sterilized cryogels were rinsed with PBS and cell culture medium. After placement on Parafilm, 60 μl of hLF suspension was seeded onto each cryogel at densities of 10,000, 20,000, 40,000 and 60,000 cells cryogel^−1^ and incubated overnight to allow cell attachment (Figure 1A). The hLF-loaded cryogels were then cultured in a humidified 5% CO_2_ incubator at 37°C with daily medium changes.

### Glucose uptake assay of hLF-loaded cryogels

Glucose uptake by hLFs cultured within cryogels was used as an indicator of cell proliferation. Culture medium was collected at designated time points, and the supernatant was obtained by centrifugation at 2,600*×g* for 10 min at 4°C. Glucose concentration in the supernatant was quantified using a glucose assay kit (Dojindo, Kumamoto, Japan). Absorbance at 450 nm was measured with a microplate reader. Glucose uptake was calculated as the difference between initial and final glucose concentrations in the medium.

### Live/dead staining of hLFs cultured within cryogels

Cell viability was evaluated using a Live/Dead Cell Staining Kit II (#PK-CA707-30002, Promocell, Germany). hLF-loaded cryogels were rinsed with PBS and incubated in the staining solution for 30 min. The kit stains live cells with calcein AM (green fluorescence) and dead cells with EthD-III (red fluorescence). After staining, the cryogels were rinsed three times with PBS, and z-stack fluorescence images were acquired using a confocal laser-scanning microscope (LSM700; Carl Zeiss, Oberkochen, Germany). Projection images were reconstructed from the z-stack using ImageJ software (National Institutes of Health, Bethesda, MD, USA).

### Preparation of microfluidic devices

The fabrication of the microfluidic devices used in this study was previously described.^[67]^ Briefly, PDMS was patterned by soft lithography and bonded to a cover glass after plasma treatment. The microchannels were coated overnight with 1 mg ml^−1^ poly-D-lysine solution (Sigma-Aldrich, St. Louis, MO, USA), rinsed twice with sterile deionized water, and dried. The fibrin-collagen hydrogel prepolymer was prepared by mixing rat-tail type-I collagen solution (2 mg ml^−1^, pH 7.4; Dow Corning, Midland, MI, USA) and fibrinogen solution (3 mg ml^−1^; F8630, Sigma-Aldrich) at a 1:9 ratio.^[68]^ Gelation was initiated by adding thrombin (5 U ml^−1^; T6634, Sigma-Aldrich), after which the prepolymer solution was promptly injected into the central hydrogel region of the device. The device was placed in a humidified box and incubated in a 5% CO_2_ incubator at 37°C for 15–20 min to complete gelation. The device comprised three parallel microchannels: the central channel was used to introduce fibrin-collagen hydrogel, while the adjacent two channels were used for seeding HUVECs and rHeps (Figure S6, Supporting Information). The rHep channel was connected to a reservoir for placing an hLF-loaded cryogel to generate a PFLG across the channel.

### Coculture of hepatocyte tissue and HUVECs in a PFLG-generating system

Perfusable hepatocyte tissues were generated in two stages using the PFLG-generating system. First, only HUVECs were cultured to construct microvascular networks (pre-angiogenesis phase). hLF-loaded cryogels were prepared as described above and rinsed with 100 μl of culture medium. A 30 μl HUVEC suspension (4×10^6^ cells ml^−1^) was introduced into the HUVEC channel of the microfluidic device (Figure 1B). The device was tilted to facilitate HUVEC attachment along the hydrogel sidewall and incubated in a 5% CO_2_ incubator for 30 min. An hLF-loaded cryogel was then inserted into the reservoir connected to the rHep channel (Figure 1B). HUVECs were cultured for 5– 7 days under PFLG, until just before the microvessels extended across the gel region.

Second, rHeps were added to generate vascularized hepatocyte tissue (vascularization phase). rHep-HUVEC mixed suspensions (30 μl, 2×10^6^ cells ml^−1^) were seeded into the rHep channel. The device was tilted to facilitate cell attachment to the hydrogel sidewall and incubated in a 5% CO_2_ incubator for 30 min. Half of the culture medium was replaced daily.

### Immunofluorescence staining

Cells were fixed in 4% paraformaldehyde for 15 min at room temperature, permeabilized with 0.1% Triton X-100 for 15 min, and blocked with BlockAce (Dainippon Pharmaceutical, Japan) for 24 h to minimize nonspecific binding. Samples were then incubated overnight at 4°C with the following primary antibodies: sheep anti-PECAM-1 (CD31) (AF806, R&D Systems, Minneapolis, MN, USA) to label HUVECs; mouse anti-multidrug resistance-associated protein 2 (MRP2) (NBP1-42349, Novus Biologicals, Centennial, CO, USA) to visualize bile canaliculi formed by rHeps; and rabbit anti-Ki67 (ab16667, Abcam, Cambridge, UK) to identify proliferating cells. After washing with PBS, appropriate secondary antibodies were applied and incubated overnight at 4°C: Alexa Fluor 555-conjugated anti-sheep IgG (Invitrogen), Alexa Fluor 647-conjugated anti-rabbit IgG (Invitrogen), or Alexa Fluor 647-conjugated anti-mouse IgG (Invitrogen). F-actin was stained with Alexa Fluor 488-phalloidin (#A12379, Invitrogen), and nuclei were counterstained with Hoechst 33342 (Invitrogen). Between each step, samples were rinsed with PBS. Z-stack images were acquired using a confocal laser-scanning microscope (LSM700; Carl Zeiss, Oberkochen, Germany), and maximum-intensity projections were reconstructed from the z-stacks using ImageJ software (National Institutes of Health, Bethesda, MD, USA).

### Quantitative analysis of vessel area and density

Microvascular networks formed during the pre-angiogenesis phase were quantified from maximum-intensity projection images of confocal z-stacks. Images were binarized using ImageJ. Vessel area fraction (%) was calculated as the percentage of vessel area within the hydrogel region of the microfluidic device. Vessel length density (µm µm^−2^) was calculated by extracting the vessel centerlines from the binarized masks (skeletonization) and dividing the total centerline length by the hydrogel region area.

### Visualization of bile canaliculi in live hepatocytes using CellTracker red CMTPX

In hepatocytes, the cell-permeant dye CellTracker Red CMTPX (C34552, Invitrogen) is metabolized by intracellular esterases to the fluorescent CMTPX, which is actively secreted into bile canaliculi via canalicular transporters.^[19]^ To visualize bile canaliculi, cells were incubated with CellTracker Red CMTPX (10 µM) in serum-free DMEM for 40 min, rinsed three times with serum-free DMEM, and then incubated in fresh medium for an additional 15 min to allow CMTPX accumulation in bile canaliculi. Z-stacks were acquired using a confocal laser scanning microscope (LSM700), and maximum-intensity projections were reconstructed from the z-stacks using ImageJ.

### Quantitative analysis of albumin secretion

Albumin secretion by rHeps in vascularized and non-vascularized hepatocyte tissues was quantified using an ELISA with the double-antibody-sandwich method. Twenty-four hours after medium change, conditioned media from each culture were collected into microtubes and centrifuged at 8,000*×g* for 10 min. Supernatants were transferred to new tubes for analysis. A 96-well plate (Nunc, Roskilde, Denmark) was coated with sheep anti-rat albumin antibody (1.0 µg well^−1^; Bethyl Laboratories, Montgomery, TX, USA). After blocking (50 mM Tris, 0.14 M NaCl, 1% bovine serum albumin, pH 8.0), samples and rat albumin standards (Bethyl Laboratories) were added and incubated for 1 h. Wells were then incubated with horseradish peroxidase-conjugated sheep anti-rat albumin antibody (2.5 ng well^−1^; Bethyl Laboratories). Tetramethylbenzidine peroxidase substrate (Kirkegaard and Perry Laboratories, Gaithersburg, MD, USA) was applied, and the reaction was stopped with 2 M H_2_SO_4_. Absorbance was read at 450 nm using a microplate reader, and albumin concentrations were interpolated from the standard curve.

### Perfusability test using fluorescent microbeads

Perfusability of vascularized hepatocyte tissues was assessed on day 5 of the vascularization phase. After immunofluorescence staining for CD31 and F-actin, samples were rinsed with PBS and mounted on an inverted fluorescence microscope (Axio Observer; Carl Zeiss) equipped with a CMOS camera (BFS-U3-13Y3M-C, Teledyne FLIR). PBS containing 1-μm fluorescent polystyrene microbeads (FluoSpheres, F8814, Invitrogen) was introduced into the inlet of the HUVEC channel of the microfluidic device. Time-lapse image sequences of bead motion within the microvessels were recorded to verify luminal continuity and perfusion in the vascularized hepatocyte tissue.

### Perfusion analysis using FITC-dextran

Functional perfusion of vascularized hepatocyte tissues was assessed on day 5 of the vascularization phase using 70-kDa FITC-dextran. After immunofluorescence staining for CD31 and F-actin, samples were rinsed with PBS and mounted on a confocal laser-scanning microscope (LSM700; Carl Zeiss). PBS containing 100 µg ml^−1^ 70-kDa FITC-dextran (Sigma-Aldrich) was introduced into the inlet of the HUVEC channel of the microfluidic device. Time-lapse fluorescence images were acquired to monitor dextran perfusion. For quantitative analysis, regions of interest (ROIs) were defined within the hepatocyte tissue (H_1_–H_5_) and hydrogel (V_1_–V_5_) domains. Fluorescence intensity in each ROI was measured to quantify dextran accumulation in hepatocyte tissue and hydrogel domains.

### Statistical analysis

Cell culture experiments were performed in at least two independent biological replicates. Data are presented as means ± SD. Group differences were analyzed using one-way analysis of variance (ANOVA), followed by Tukey’s post hoc test for multiple comparisons. Statistical significance was set at *P <* 0.05.

## Supporting Information

Supporting Information is available from the Wiley Online Library or from the author.

## Acknowledgements

This study was supported by the Japan Society for the Promotion of Science (grant number: 25K03439).

## Data Availability Statement

The data that support the findings of this study are available from the corresponding author upon reasonable request.

Received: ((will be filled in by the editorial staff))

Revised: ((will be filled in by the editorial staff))

Published online: ((will be filled in by the editorial staff))

## Supporting Information

**Figure S1.**
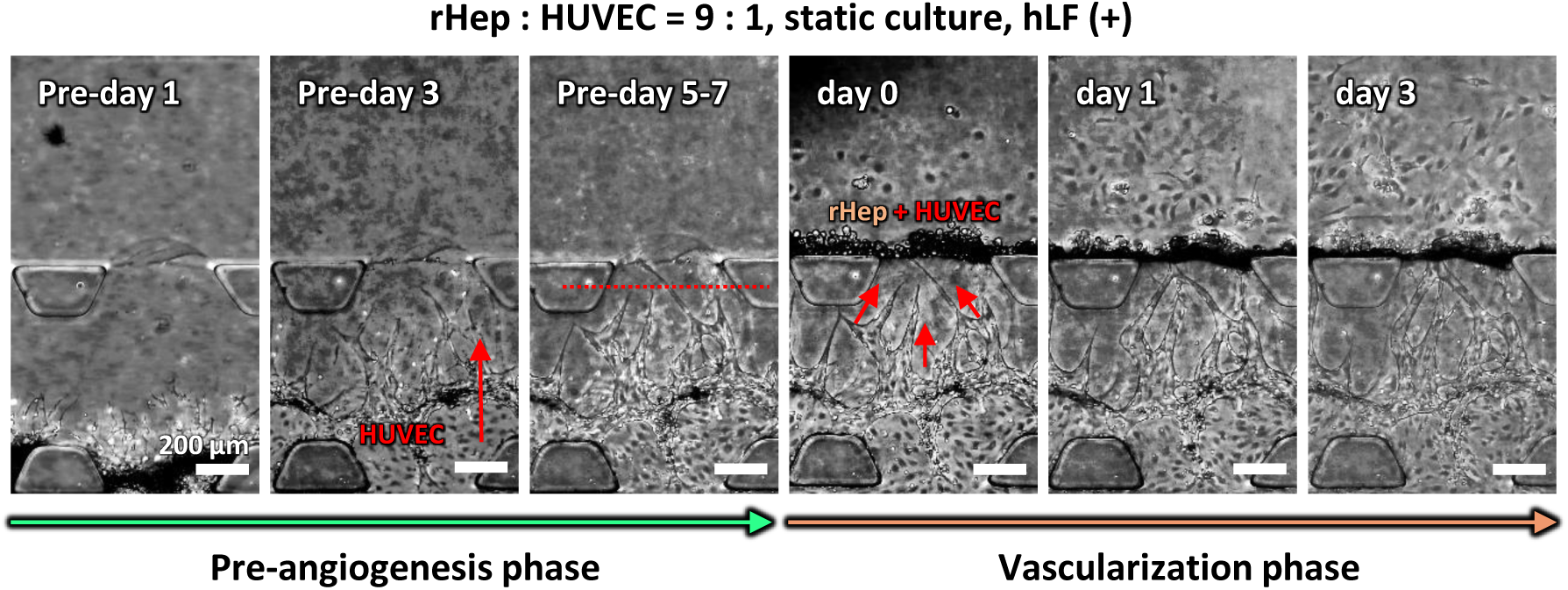
The process of pre-angiogenesis and subsequent vascularization of hepatocyte tissue. Corresponding phase-contrast images showing the progression of pre-angiogenesis and subsequent vascularization of hepatocyte tissue. During the pre-angiogenesis phase, HUVECs formed angiogenic sprouts under the PFLG generated by hLF cryogels. In the vascularization phase, microvessels penetrated into the hepatocyte tissue. Scale bars, 200 µm.

**Figure S2.**
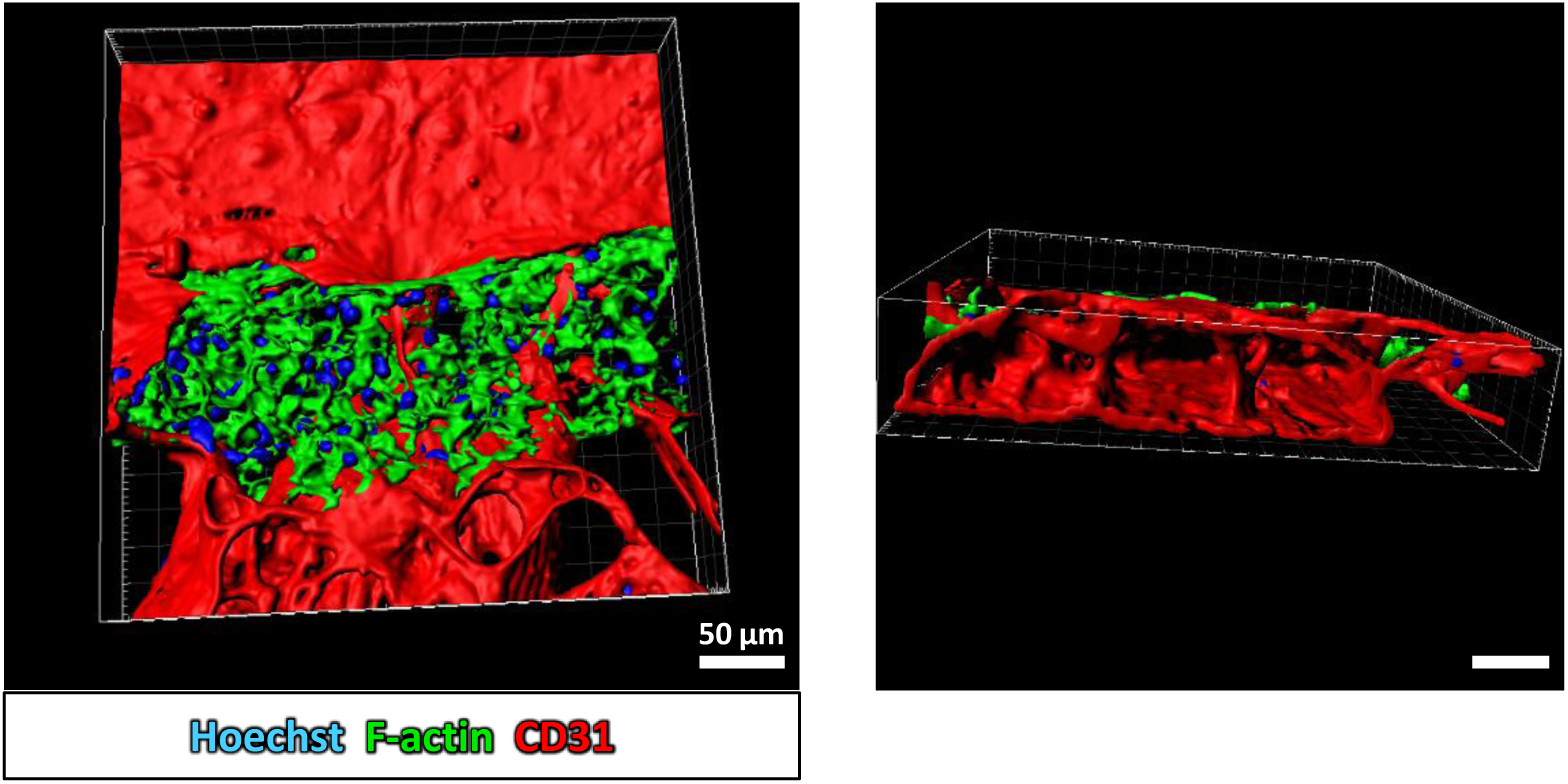
Luminal structure of microvessels in vascularized hepatocyte tissue. Three-dimensional reconstruction images of vascularized hepatocyte tissue, demonstrating luminal microvessels penetrating into the hepatocyte tissue and recapitulating the architecture of hepatic sinusoids. Blue: cell nuclei (Hoechst 33342), green: F-actin marking hepatocyte tissue (Alexa Fluor 488-phalloidin), red: microvessels (CD31). Scale bars, 50 µm.

**Figure S3.**
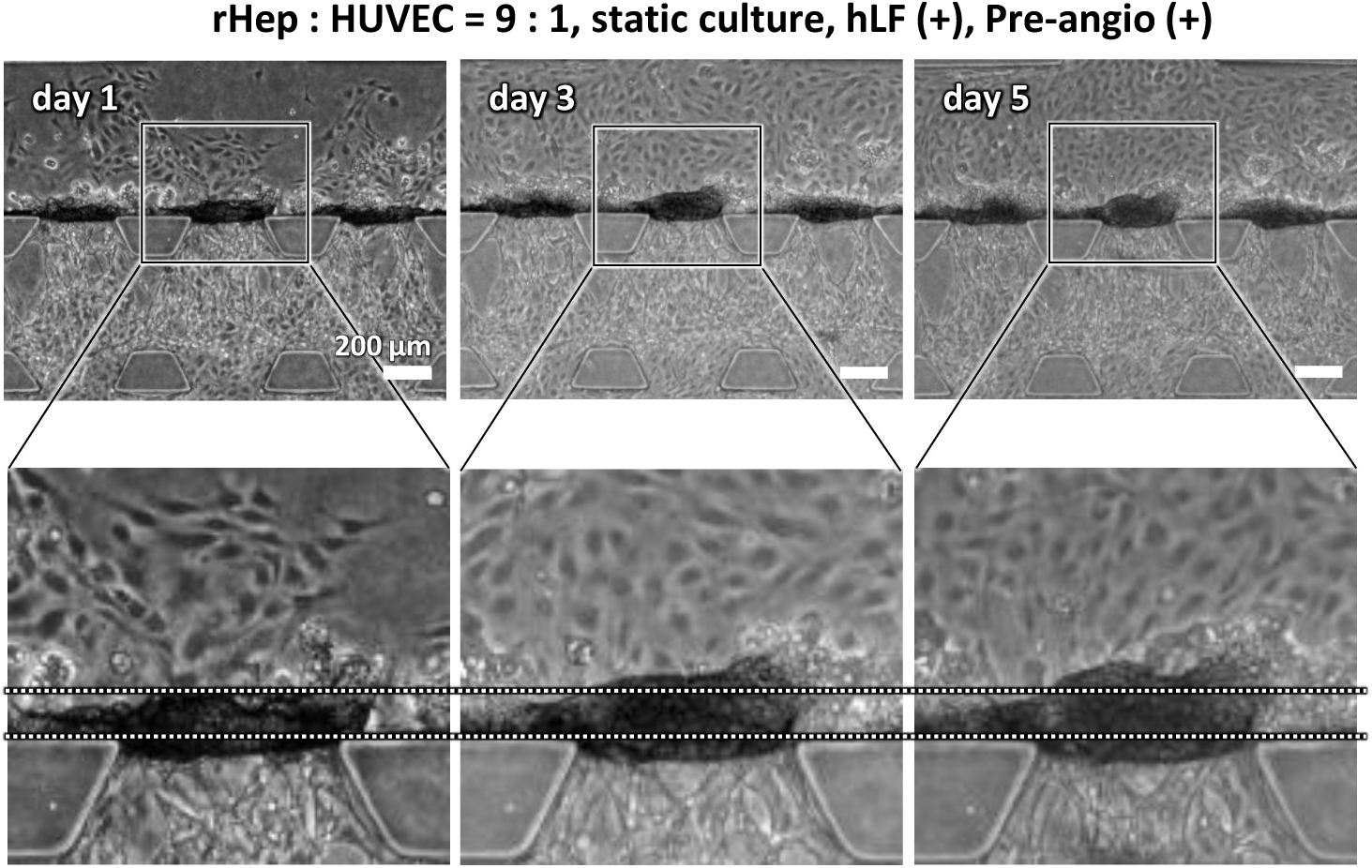
Growth of vascularized hepatocyte tissue. Phase-contrast images showing progressive growth of vascularized hepatocyte tissue from day 1 to day 5. Scale bars, 200 µm.

**Figure S4.**
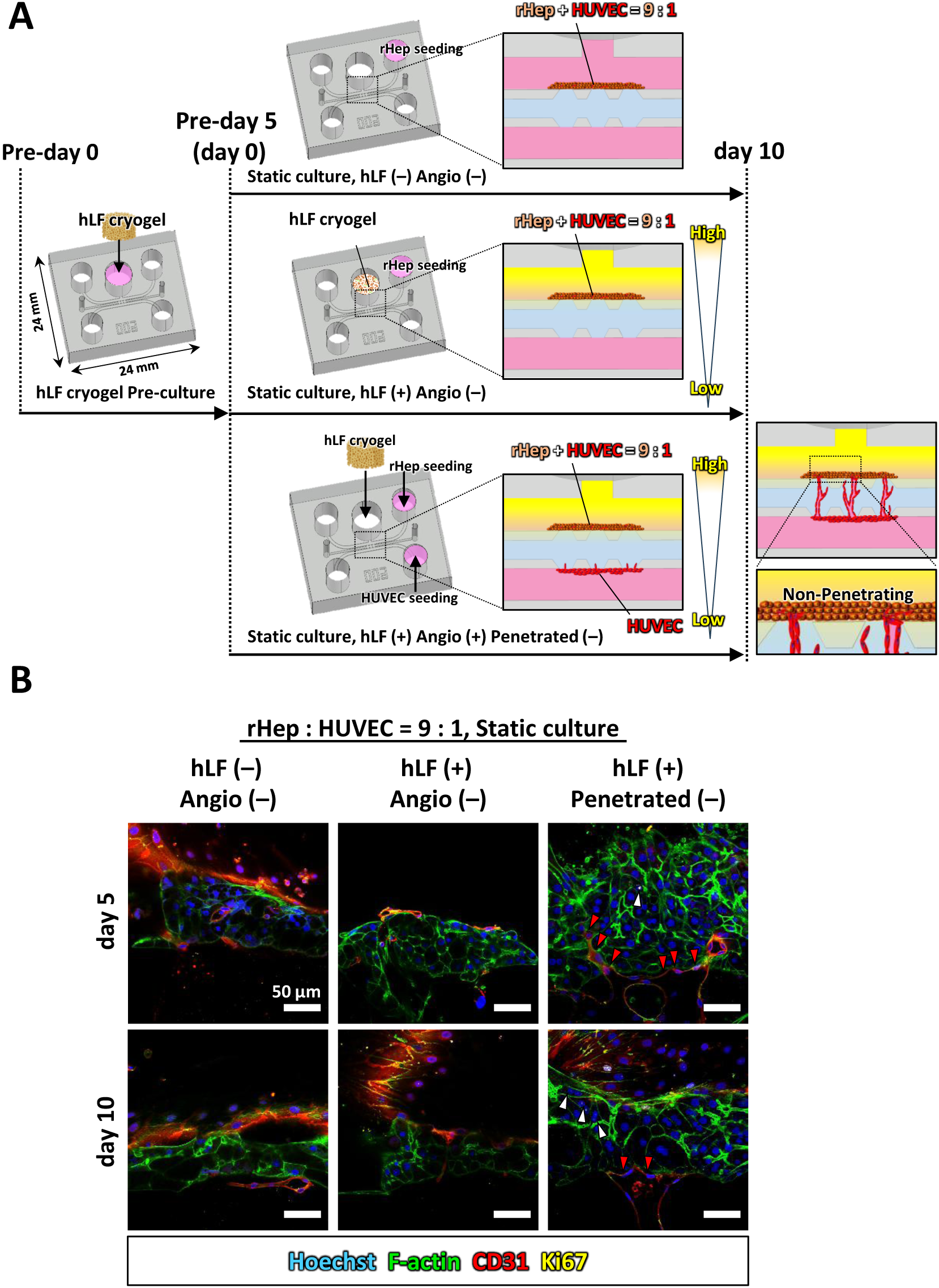
Hepatocyte proliferation in non-vascularized hepatocyte tissue. A: Experimental design and timeline showing three different culture conditions: hLF (–) Angio (–), hLF (+) Angio (–), and hLF (+) Angio (+) Penetartraed (–). B: Immunofluorescence images of cell nuclei (blue: Hoechst 33342), F-actin (green: Alexa Fluor 488-phalloidin), HUVECs (red: CD31) and proliferating cell nuclei (yellow: Ki67). White arrowheads indicate Ki67-positive hepatocytes. Red arrowheads indicate non-penetrating microvessels at the boundary of hepatocyte tissue. Scale bars, 50 µm.

**Figure S5.**
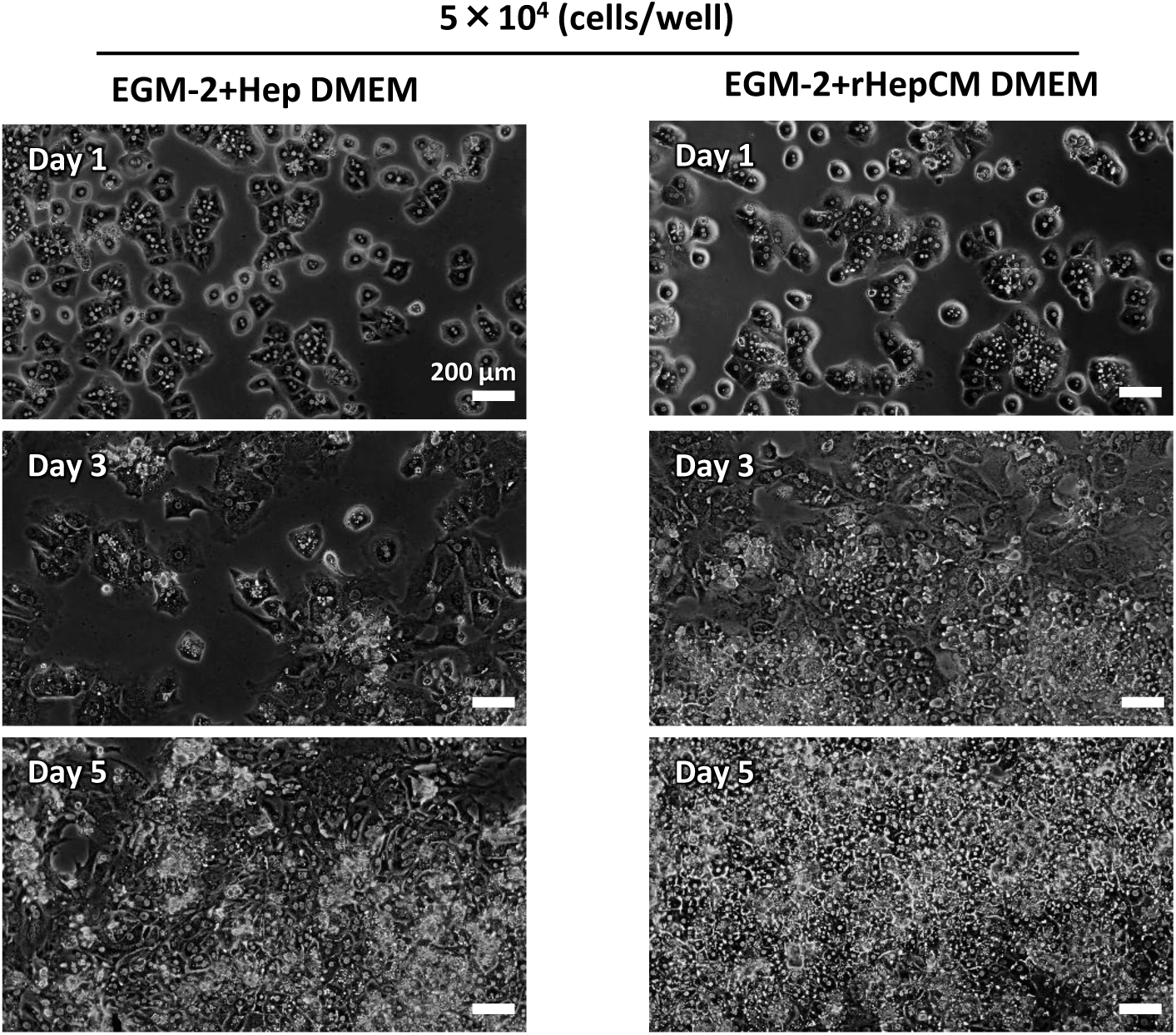
Morphological response of rHeps depending on culture media. Phase-contrast images of rHeps in 2D culture comparing EGM-2 + Hep DMEM and EGM-2 + rHep CM over 5 days. Initial seeding density: 5 × 10⁴ cells well^−1^. Scale bars, 200 µm.

**Figure S6.**
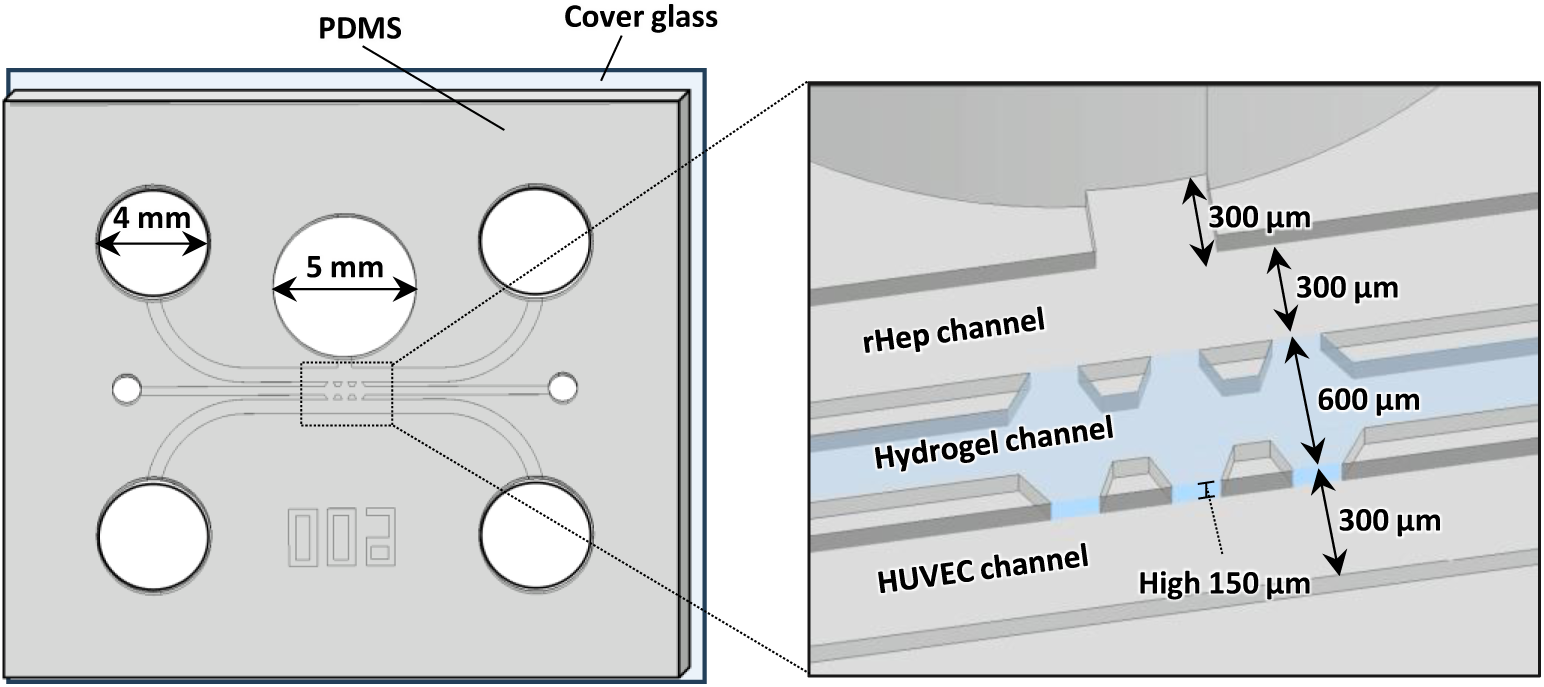
Schematic illustrations of microfluidic device design.

Video S1. Luminal structure of microvessels in vascularized hepatocyte tissue. Movie corresponding to Supplementary Fig. 3.

Video S2. Perfusion of FITC-dextran solution through vascularized hepatocyte tissue. Time-lapse movie was captured using a confocal microscope showing the perfusion of 70 kDa FITC-dextran solution through vascularized hepatocyte tissue. Green: FITC-dextran and F-actin (Alexa Fluor 488-phallidin), red: microvessels (CD31).

Videos S3. and S4. Fluorescently labeled microbeads perfusion. Time-lapse movies showing the perfusion of fluorescently labeled microbeads (diameter: 1 µm). Sequential fluorescent images were captured using a fluorescence microscope equipped with a high-speed camera.

Video S5. Real-time CMTPX excretion by polarized hepatocytes. Movie showing cytoplasmic CMTPX being excreted into bile canaliculi (red: CMTPX

